# LinkLlama: Enabling Large Language Model for Chemically Reasonable Linker Design

**DOI:** 10.64898/2026.04.15.718690

**Authors:** Kunyang Sun, Yingze Wang, Justin Purnomo, Joseph M. Cavanagh, Giovanni Battista Alteri, Teresa Head-Gordon

## Abstract

Fragment-based drug discovery (FBDD) relies heavily on the design of chemically viable linkers to connect fragments binding to different pocket regions into potent lead molecules. While recent generative models have advanced spatial fragment linking, they frequently produce linkers characterized by high torsional strain and non-drug-like motifs. In this work, we present LinkLlama, a fine-tuned Meta Llama 3 model that bridges the gap between text-based generation and 3D spatial awareness. By accepting natural language prompts that specify geometric constraints, such as distances and angles, alongside physicochemical targets like Lipinski’s rules and rotatable bond limits, LinkLlama generates highly tailored molecules for the input fragments. Leveraging the inherent chemical grammar captured through supervised fine-tuning on a curated corpus of drug-like molecules from ChEMBL, the model prioritizes chemical validity without requiring complex reinforcement learning loops. Benchmarking on the ZINC and HiQBind datasets demonstrates that LinkLlama maintains competitive geometric fidelity compared to strictly 3D-aware models while achieving a two-fold increase in the proportion of chemically reasonable designs. This rising success rate, jumping from 35% to over 80%, is defined by strict adherence to comprehensive structural filters including PAINS, non-drug-like chemical patterns and complex ring systems. We further illustrate the model’s versatility through prospective case studies in novel small-molecule scaffold hopping and PROTAC linker design, validated via molecular docking and molecular dynamics simulations against known crystal poses. Ultimately, LinkLlama demonstrates that large language models can overcome the structural pitfalls of purely 3D-generative methods, offering a highly controllable and chemically robust framework to accelerate linker design and drug discovery in general.

## Introduction

Fragment-based drug discovery (FBDD) has established itself as a cornerstone of modern medicinal chemistry^1–6^. By identifying low-molecular-weight “hits” that bind to specific protein sub-pockets, FBDD provides an efficient starting point for exploring chemical space. However the central challenge in FBDD is the transition from individual fragments to lead-like molecules, a process that requires the design of optimal linker chemistry^7,8^. Effective linker design is essential for connecting fragments from X-ray soaking experiments, performing scaffold hopping to replace problematic functional groups, and executing general hit expansion strategies^9^. However, the design of a linker is a complex multi-objective and constrained optimization task. Beyond simply connecting fragments, a chemically reasonable linker must ensure that the fragments maintain their original binding orientations. Furthermore, the resulting compound should exhibit low internal strain and possess drug-like properties while avoiding synthetically inaccessible motifs or undesired functional groups. With the advancement of machine learning for drug discovery, several models have been developed to automate this process, broadly categorized into 2D and 3D approaches. 2D models such as DeLinker ^10^ and Link-INVENT^11^ have demonstrated success in generating novel scaffolds. DeLinker utilizes an iterative atom-growth strategy, whereas Link-INVENT employs a reaction-based fragmentation and masked-encoder approach that can be integrated with reinforcement learning (RL) to optimize specific molecular properties. Conversely, 3D-aware models like DELETE^12^ and DiffLinker ^13^ leverage Graph Neural Networks (GNNs) and diffusion models to propose linkers directly within a three-dimensional coordinate space, making them suitable methods for pocket-aware linker generation.

Despite these advancements, existing methods face significant limitations. The 2D models often require extensive RL or heavy post-hoc filtering to achieve the conditional sampling that satisfies medicinal chemistry standards. While 3D models are spatially aware, they frequently produce structures with geometric errors, such as unrealistic bond lengths or highly strained torsions, which may force fragments out of their bioactive conformations. Furthermore, standard evaluation metrics like the Quantitative Estimate of Drug-likeness (QED)^14^ and Synthetic Accessibility (SA) scores^15^ can be misleading. These metrics are often dominated by the properties of the initial fragments and fail to provide granular information regarding the quality of the generated linker itself. Furthermore, as pointed outed by Walters^16^, current models suffer greatly in generating molecules with stable drug-like chemistry, often rendering these models futile in real drug discovery deployment.

The recent success of Large Language Models (LLMs) in chemistry has opened new avenues for molecular and linker design to address these challenges. As foundation models^17^, LLMs inherently possess broad chemical knowledge. Recent studies demonstrate that this representational capacity can be efficiently adapted to specific chemical tasks through supervised fine-tuning (SFT)^18–22^. Several recent works exemplify this paradigm: SmileyLlama^23^ showed that fine-tuning an open-weight LLM produces a Chemical Language Model (CLM) performing at or above the level of models trained exclusively on SMILES strings, utilizing natural language prompts to enable direct conditioning on molecular properties. Similarly, SynLlama^24^ demonstrated that LLM fine-tuning can overcome synthesizability bottlenecks by constraining molecule generation to a manifold of commercially available building blocks and established chemical reactions. Ultimately, by coupling the deep internal chemical priors of an LLM with carefully curated fine-tuning datasets, complex chemical design problems can be effectively steered and resolved using natural language, typically with much less data and without RL retraining.

In this work, we present LinkLlama, a framework designed to bridge the gap between computational linker generation and practical medicinal chemistry. Our primary focus is on chemical reasoning that ensures the model consistently generating highly realistic molecules that chemists can readily act upon. We achieve this practical utility through an “alignment-by-design” approach that enables reinforcement learning (RL)-free conditional sampling. Rather than relying on computationally expensive RL cycles or complex reward-function engineering to optimize multiple objectives ^11,25,26^, users can dynamically steer LinkLlama via natural language prompts to satisfy precise structural and physicochemical constraints. This level of control is highlighted in our case analyses, where the model successfully navigates diverse design challenges, from standard scaffold hopping to satisfying the demanding architectural requirements of PROTAC linkers, while maintaining the vital spatial integrity of fragment hits. Ultimately, by combining robust chemical constraints with intuitive natural language guidance, LinkLlama emerges as a versatile and powerful tool for property-driven linker design.

## Methodology

### Input Preparation

#### ChEMBL Data Processing

To obtain training data that covers a diverse, drug-like chemical space, we use ChEMBL36^27^ due to its breadth relative to earlier linker-generation work based on ZINC^10^. We apply a cleaning procedure of this database, which filters out molecules containing low-frequency atoms, exhibiting complex chemistry (such as low carbon fractions or complex ring systems), possessing excessive molecular weights, or being too small for fragmentation. Ultimately, approximately 93.0% of the ChEMBL36 molecules survive this culling, yielding a valid set of 2,665,082 molecules. The full cleaning steps are detailed in Supplementary Information S1. Distributions comparing ChEMBL to other libraries (including PROTAC-relevant chemical space) are shown in Supplementary Figure S3.1.

#### Molecular Fragmentation

Once cleaned, the remaining ChEMBL molecules are fragmented into fragment-linker-fragment triplets following RDKit^28^ matched molecular pair analysis. We perform exactly two cuts per molecule to generate these triplets, restricting the cuts strictly to acyclic single bonds incident to neutral sp^3^ carbons. Because a single molecule can be fragmented in multiple ways, we apply specific criteria to filter the resulting decompositions. We retain cuts only if both the linker and fragments meet minimum heavy-atom counts, the linker possesses a topological path length of at least two between attachment points, and the linker is not larger than the smallest terminal fragment. Full details are provided in Supplementary Information S1. From the cleaned ChEMBL36 set, this process yields 8,303,935 fragment-linker triplets with 256,135 unique linkers.

#### Assessment of Chemically Reasonable Molecules

We construct a rigorous analysis for both the designed linker and the complete molecule, utilizing two evaluation benchmarks from Walters and three additional ”chemical reasonability” rules from iMiner^29^. We distinguish between properties specific to the isolated linker and those applied to the whole molecule. For the linker itself, we check for overly complex bridgehead ring structures or uncommon ring systems (defined as occurring fewer than 100 times in the entire ChEMBL36 database) ^30,31^. If a linker contains either motif, it is classified as “unreasonable.” At the whole-molecule level, we apply a set of iMiner undesirable SMARTS^32^ patterns that document common generative model errors, alongside PAINS^33^ and Brenk filters^34^ to identify and reject non-drug-like or problematic chemical motifs. Overall, a final molecule is classified as “reasonable” only if it passes all five metrics. Detailed implementations are provided in Supplementary Information S1.

#### Supervised Fine-tuning and Inference from LinkLlama

To create the LinkLlama model, we establish data generation protocols to format the cleaned input preparation of the ChEMBL dataset into a supervised fine-tuning (SFT) corpus. We frame the task as an instruction-following completion in Alpaca-style JSONL format. Each record provides a fixed instruction, a natural-language input that encodes the fragment pair and optional constraints, and a structured output. As illustrated in Figure 1, the input always states the SMILES of both terminal fragments and their distance (in Å) and angle (in degrees). The distance is captured by the distance between the two anchor atoms, and the angle is calculated between the two unit vectors from each anchor atom pointing towards their connected dummy atoms. Additional linker-type and physicochemical band constraints (linker rotatable bonds, linker heavy atoms, Hbond donors/acceptors, molecular weight, log *P*, TPSA) are included stochastically, each with an independent 50% chance during generation. The overall reasonable/unreasonable label is never dropped and is derived from the five reasonability checks as defined in the previous section. The complete template, all permissible input constraint strings, and the training randomization rules are documented in Supplementary Information S1.

**Figure 1:**
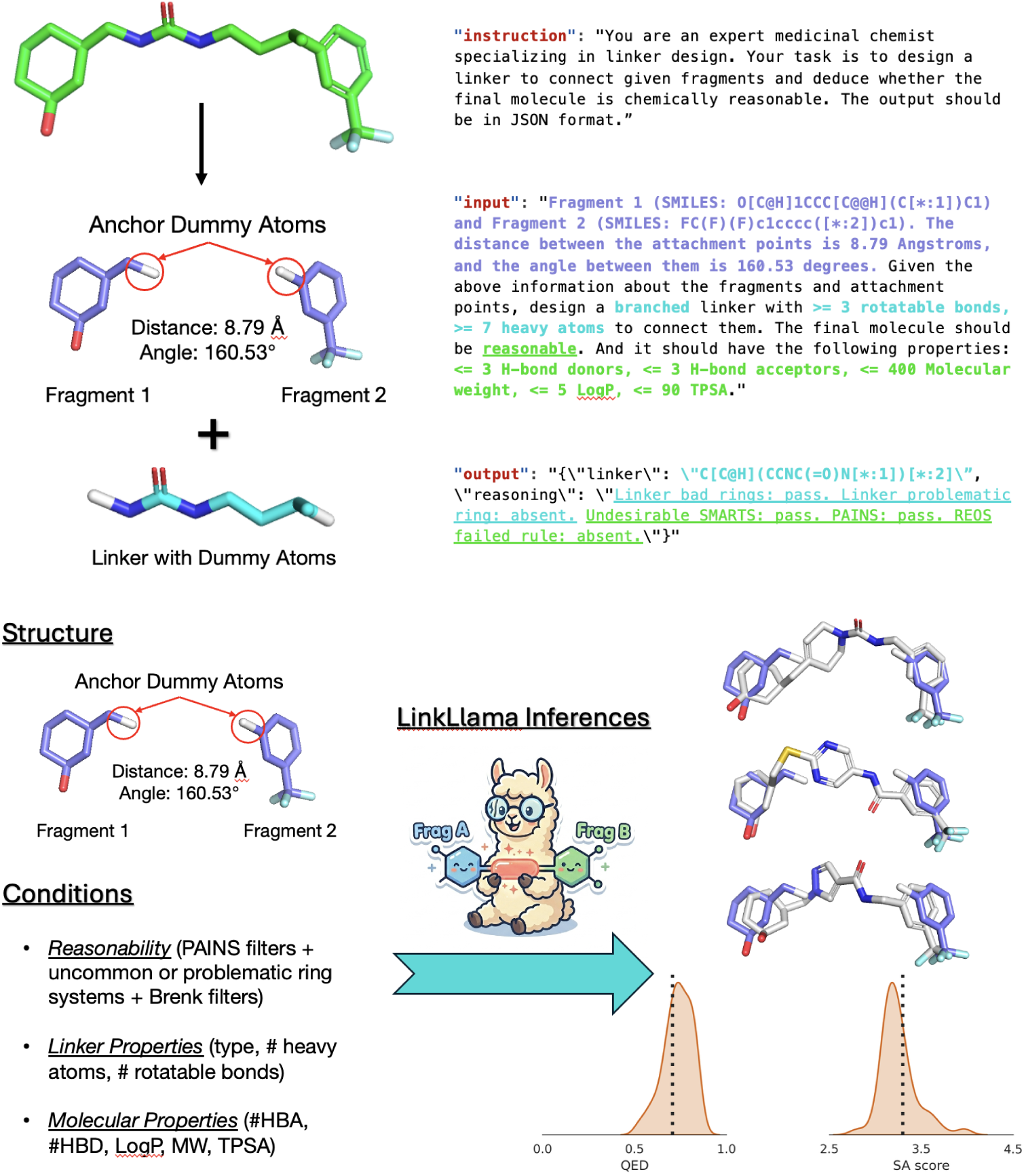
(Top). Schematic representation of the LinkLlama data curation pipeline, illustrating the transition from raw ChEMBL molecules to the filtered SFT corpus via fragmentation and chemical reasonability checks + the prompt construction and result processing. (Bottom). LinkLlama inference process with two parts of input specified in the SFT corpus along with 3 example generated molecules overlayed on crystal fragment poses and the distribution fo QED and SA score for the 100 LinkLlama-generated whole molecules. The LinkLlama cartoon image is generated by Google Gemini.

The model’s output is structured to predict not only the intervening linker but also to explicitly evaluate the chemical reasonability of the training structure. The expected response is a JSON object with the linker SMILES and a reasoning string that records the pass/fail status for the same chemical filters summarized above. By jointly modeling linker generation and this reasonability trace, LinkLlama learns to condition its generative space on medicinal chemistry heuristics.

LinkLlama is developed by fine-tuning the Meta Llama-3.2-1B-Instruct checkpoint on a subset of the prepared ChEMBL36 data filtered to minimize the over-representation of frequent linkers. This final subset contains 1,526,169 training examples. Fine-tuning is executed on a single node with 4 *×* A100 GPUs in approximately 6.5 hours. Full training details are listed in Supplementary Information S1.

During inference, prompts use the same sentence template, but the fragment geometry is always taken from the input data, while linker-type bands, molecule-property bands, and the requested reasonable/unreasonable condition are set explicitly in a YAML configuration. The model returns a JSON object with a linker SMILES and a reasoning string, where the linker SMILES string is then merged with the input fragments into the full molecule by aligning the anchor dummy atoms on the predicted linker and input fragments.

#### Hyperparameter Optimization and Model Selection

We first fine-tune a baseline model on two million randomly sampled fragment-linker pairs drawn from the full processed ChEMBL36 pool and assess its performance. In the initial pilot run, our primary focus is the validity, uniqueness, and novelty of the generated linkers produced by this model on the external ZINC-hard dataset used in the Results section. Supplementary linker-frequency diagnostics (Supplementary Figure S3.2) reveal a severe long tail in the original ChEMBL data distribution, where a small set of simple yet common linkers, such as amide bonds, dominates the training counts. In practice, this checkpoint achieves high validity on the ZINC-hard dataset but relatively low uniqueness among valid generations and modest novelty with respect to the training linker vocabulary. As shown in Figure 2, this trend confirms the potential memorization of the high-frequency head of the linker distribution rather than the exploration of diverse bridges. To counteract this collapse in uniqueness, we curate two rebalanced training pools. First, the Cap50 corpus applies a strict per-linker frequency ceiling of 50 counts. Specifically, if a linker has more than 50 occurrences, we select 50 diverse samples based on their molecular properties and drop all other entries. This hard cap flattens the occurrence histogram and increases effective linker diversity at the cost of modest shifts toward heavier, more flexible linkers (Supplementary Figure S3.3), yielding the Diverse 1.5M split with 1,536,170 training samples. In addition, the Hybrid corpus employs a softer capping rule that preserves more of the natural rank-frequency ordering while still suppressing the most extreme repeats. In this approach, we apply the same diversity-based sampling for down-selection: linkers with fewer than 50 rows remain unchanged, those with between 50 and 500 rows are reduced by 10%, and those with more than 500 rows are hard-capped at 450 (i.e., 500 - 500 *×* 10%). This procedure results in the **Diverse 2.9M** split containing 2,817,412 training samples. We train separate Llama-3.2-1B LoRA models on the Random 2M (unbalanced draw from the full pool), Diverse 1.5M (Cap50), and Diverse 2.9M (Hybrid) splits and compare them under identical evaluation protocols.

**Figure 2:**
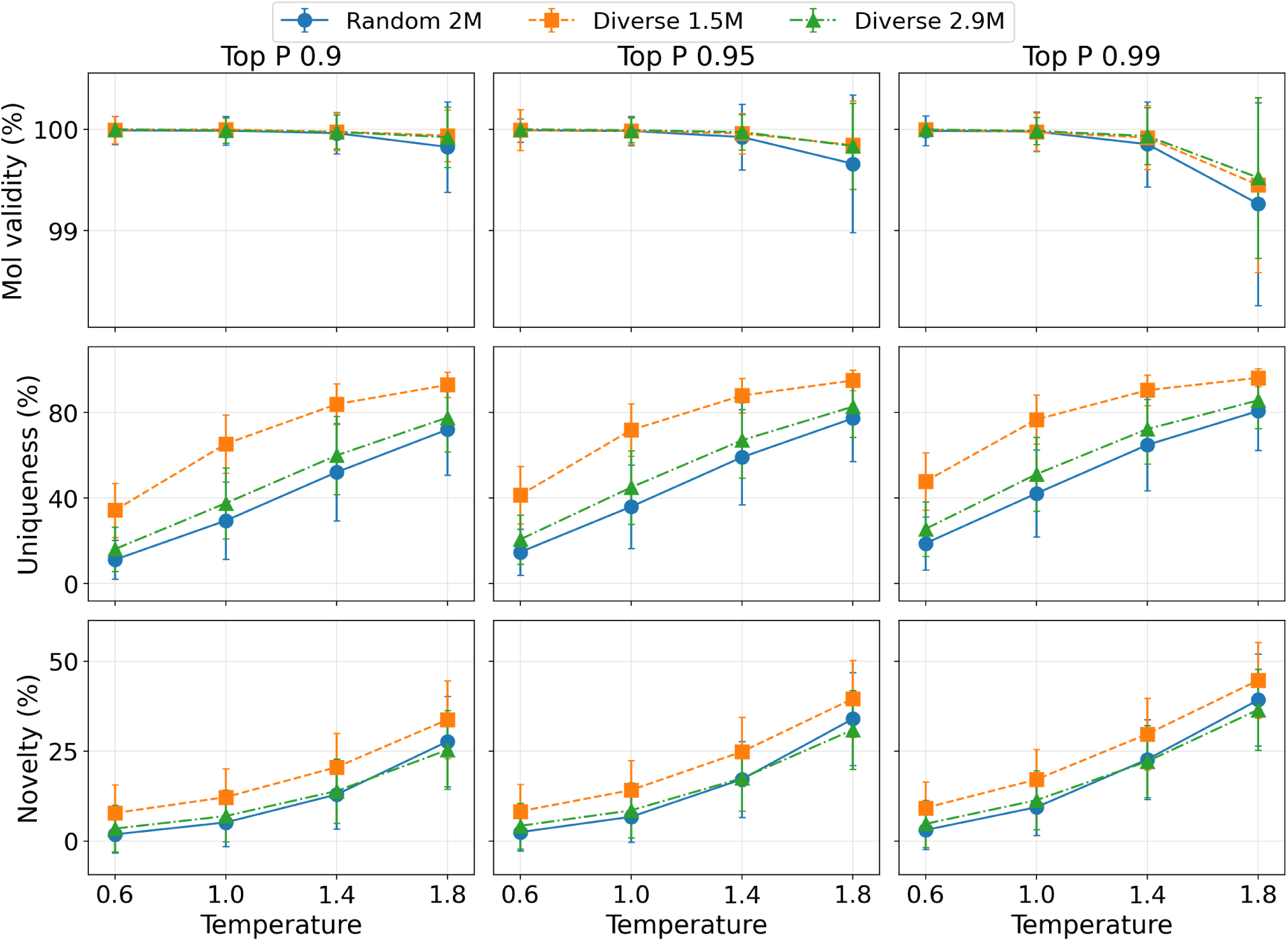
Model hyperparameter searching performances of unconditional generation on the ZINC-hard dataset. The legend denotes the three evaluated models trained on different data splits: **Random 2M** (the long-tailed, unbalanced pool), **Diverse 1.5M** (the strict Cap50 rebalanced pool), and **Diverse 2.9M** (the hybrid-capped pool). Individual data points represent temperature settings (*T* ∈ {0.6, 1.0, 1.4, 1.8}), and the columns correspond to Top-P sampling values (top-*p* ∈ {0.9, 0.95, 0.99}). The rows display mean molecule validity (%), uniqueness among valid samples (%), and the novelty of valid linkers compared to the training linker set (%), with shared *y*-axes within each row. Error bars indicate the dispersion across instances and aggregated runs.

In regards hyperparameters, we sweep temperature (*T*) and Top-P (top-*p*) as summarized in Figure 2. The Diverse 1.5M (Cap50) checkpoint consistently offers the best compromise among validity, uniqueness, and novelty on this grid. As shown in Supplementary Figure S3.5, grids on another set we use in the Results section, the ZINC-random set, also uncovers the same overall trend. Therefore, we use this model with *T* = 1.4 and top-*p* = 0.99 everywhere below.

## Results

### Standard Benchmarking on ZINC

For the first set of generation tasks, we use the ZINC database^35^, which is orthogonal to our ChEMBL training set. Although LinkLlama is trained on ChEMBL, ZINC offers a screening-library-like distribution distinct from the training corpus. Furthermore, it serves as the standard test evaluation setting in prior works, ensuring direct comparability of LinkLlama with established 2D and 3D baseline models. To perform a thorough evaluation, we construct a random 1k subset of fragment pairs to represent typical generation difficulty, alongside a hard 1k subset. In the hard 1k subset, the reference molecule is precomputed to pass all five chemistry checks (ensuring chemical reasonability), and the reference linker contains either more than three rotatable bonds or more than seven heavy atoms. This hard test stresses the models’ ability to design long or flexible bridges following the structural rules defined in the Methods section. After selecting the SMILES strings for each subset, we use RDKit to generate a conformer for each molecule and apply the MMFF94 force field to relax the energy. We then save the fragments into SDF files and calculate their structural input information for LinkLlama inferences.

During model inference, we evaluate the unconditional generation capabilities of LinkLlama to ensure a fair comparison. Under this constraint, all optional input keywords are omitted, and only the reasonability condition is enabled. Therefore, the input prompt is structured as follows:

FRAGMENT INFO Given the above information about the fragments and attachment points, design a linker to connect them. The final molecule should be reasonable.

Using this prompt, we compare LinkLlama against DeLinker (representing 2D graph generation baseline) and DiffLinker (representing 3D diffusion baseline). The input fragment SDF files are prepared for baseline method inferences.

To comprehensively assess generation quality, we measure a suite of standard metrics. *Validity* tracks the percentage of generated candidates per test instance that yield a well-formed product. Specifically, the structure must pass successfully in RDKit, carry compatible charges with the input fragments, and match the intended anchor connectivity. Among these valid outputs, *uniqueness* calculates the fraction of distinct canonicalized full-molecule SMILES, collapsing duplicate stereoisomer or tautomer representations. For LinkLlama, we additionally report *novelty*, which measures the fraction of valid, distinct linker graphs absent from the ChEMBL-derived training data inventory; novelty values for the baseline methods are extracted from Igashov et al. ^13^. To evaluate general drug-likeness and synthesizability, we compute RDKit’s quantitative estimate of drug-likeness (*QED*, bounded in [0, 1]) and synthetic accessibility score (*SA*, where higher values indicate more difficult syntheses). Finally, *reasonability* represents the percentage of valid molecules that pass our composite medicinal-chemistry screen, which encompasses linker-centric ring rules alongside whole-molecule PAINS, Brenk, and related structural alerts.

Table 1 reports these benchmark metrics. On the random split, LinkLlama nearly saturates validity while maintaining competitive uniqueness, whereas DiffLinker shows the largest bootstrap uncertainty and the lowest validity and uniqueness among the three methods. Although the novelty score of LinkLlama appears to fall short compared to the baselines, novelty must be balanced with chemical realism, because the generated linker and overall chemistry should still be synthetically attainable and structurally sound. Therefore, the novelty score should be interpreted alongside the reasonability percentage. LinkLlama effectively doubles the reasonability rate relative to DiffLinker and substantially outperforms DeLinker, confirming that the model successfully conditions its generative space on chemistry that passes the five reasonability checks. QED and SA sit in a similar band across all baselines, with LinkLlama showing marginal improvements. On the hard split, the performance gap becomes even more pronounced. While DiffLinker’s validity rises relative to the random subset, its uniqueness remains the lowest. Conversely, LinkLlama’s uniqueness slightly exceeds DeLinker’s, its novelty increases, and its reasonability advantage widens dramatically, nearly doubling the success rates of the baseline methods. This observation indicates that in these harder, more drug-discovery-relevant cases, LinkLlama produces more sensible designs that medicinal chemists can readily utilize.

**Table 1:** Generative performance comparison on the ZINC random 1k and ZINC hard 1k benchmarks. Validity, uniqueness, novelty, and reasonability are reported as percentages, while QED and SA are dimensionless. For the percentage columns, values represent the instance-aggregated mean ± the bootstrap standard deviation of that mean (using 2000 bootstrap resamples). Reasonability represents the average per-instance pass rate across all instances. QED and SA are reported as the instance-mean ± standard deviation across instances.

| Split | Model | Validity | Uniqueness | Novelty | QED | SA | Reasonability |
| --- | --- | --- | --- | --- | --- | --- | --- |
| Random 1k | DeLinker | 96.0±0.2% | 86.4±0.1% | 47.1%* | 0.63±0.11 | 3.39±0.63 | 44.2±0.8% |
| Random 1k | DiffLinker | 89.8±0.8% | 66.1±0.8% | 47.7%* | 0.65±0.11 | 3.42±0.74 | 25.1±0.8% |
| Random 1k | LinkLlama | 99.9±0.0% | 87.7±0.3% | 25.8±0.3% | 0.68±0.13 | 3.30±0.62 | 73.1±1.1% |
| Hard 1k | DeLinker | 89.7±0.2% | 88.3±0.2% | 47.1%* | 0.64±0.10 | 3.16±0.58 | 43.4±0.7% |
| Hard 1k | DiffLinker | 95.6±0.4% | 64.1±0.8% | 47.7%* | 0.63±0.09 | 3.05±0.65 | 31.0±0.6% |
| Hard 1k | LinkLlama | 99.9±0.0% | 90.4±0.2% | 29.7±0.3% | 0.68±0.12 | 2.99±0.55 | 87.4±0.5% |

\* Data extracted from DiffLinker Table 1. Baseline models were trained on ZINC dataset which has less unique linkers.

Beyond 2D graph validity, successful linker design requires physically realistic 3D geometries. Figure 3 summarizes 3D-aware strain and pose agreement on the ZINC datasets. Because DeLinker and LinkLlama generate SMILES strings as their final output, we assess their pose RMSD using the same generation procedure employed to create the ZINC input data: RDKit generates an initial conformer, which is subsequently relaxed via the MMFF94 force field. We quantify pose agreement using the mean and best fragment RMSD, which compare the heavy-atom positions of the generated ligand to the ZINC decoy reference after alignment on the fixed fragments (lower values indicate a closer match to the reference pose). To evaluate physical realism, we calculate the mean MMFF Δ*E* after relaxing each pose. This metric is defined as the difference between the MMFF94 energy of the proposed full molecule (after a short force-field relaxation) and the reference ligand, serving as a proxy for internal strain where larger positive values indicate energetically unfavorable proposals. While the RMSD violin plots show that all methods achieve roughly overlapping pose agreement, their strain distributions differ significantly. DiffLinker’s Δ*E* distributions extend to much higher strain limits, suggesting frequent generation of distorted or sterically clashing geometries. In contrast, LinkLlama and DeLinker consistently produces low-strain structures, indicating that it naturally favors physically viable conformations even without explicit 3D spatial diffusion coordinates used by DiffLinker. Overall, on the ZINC test sets, LinkLlama generates diverse, unique candidates with competitive 1D and 3D assessment metrics, while yielding a significantly larger population of chemically reasonable structures.

**Figure 3:**
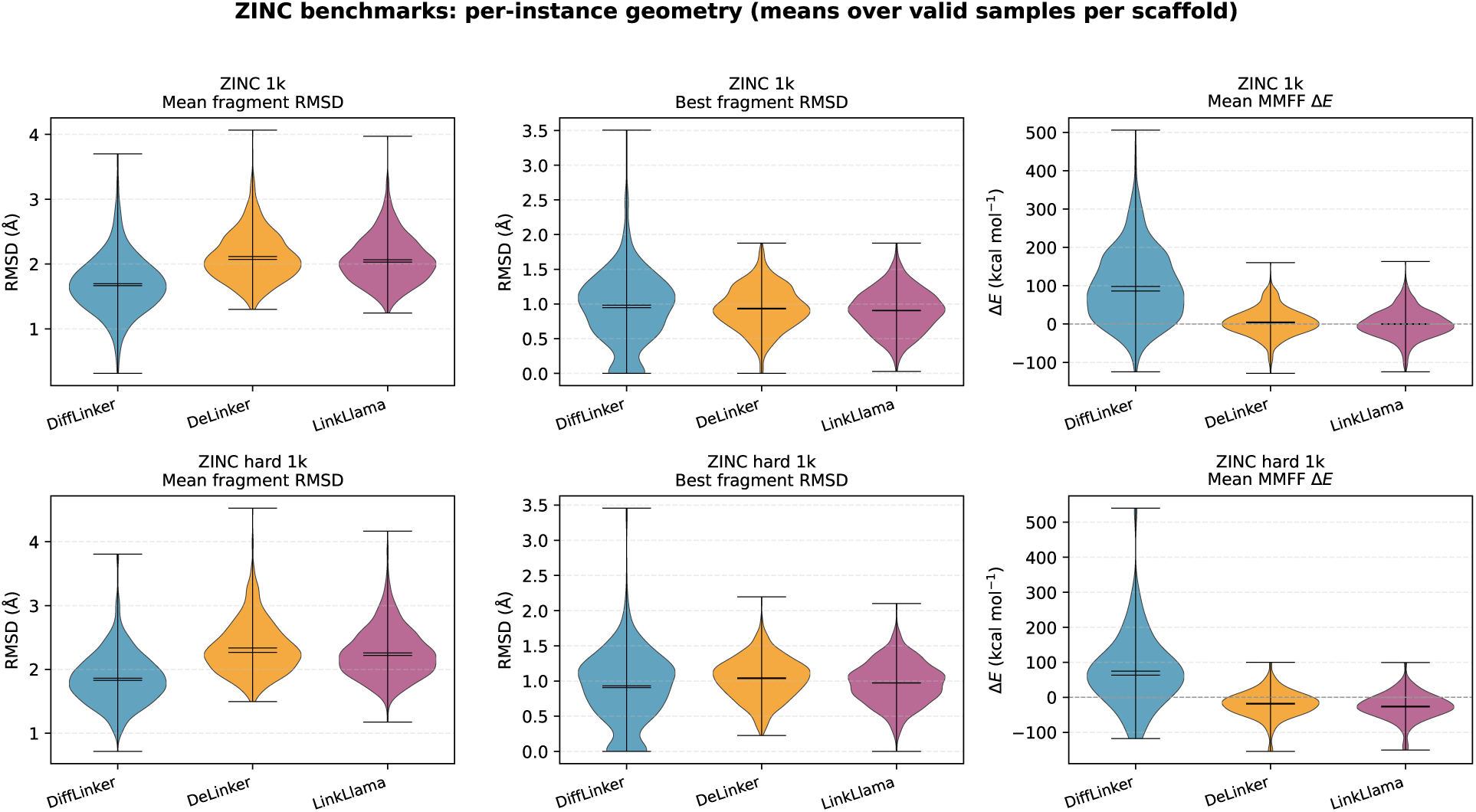
Assessment of 3D geometric realism and internal strain for generated molecules. The violin plots display the per-instance distributions for mean fragment RMSD, best fragment RMSD, and mean MMFF Δ*E* (kcal mol^−1^) across the ZINC random (top row) and hard (bottom row) 1k datasets. The three columns correspond to these distinct geometry metrics. The distributions compare the generative outputs of DiffLinker, DeLinker, and LinkLlama, highlighting differences in pose agreement with the reference and the propensity of each method to generate high-strain, energetically unfavorable configurations.

### Prompt-Conditioned Generation on ZINC Hard

Having evaluated unconditional generation, we design the next experiment to assess distribution control over the generated linkers. Specifically, we task the model with generating not only valid and reasonable linkers but also complete molecules whose physicochemical properties and linker topologies match specific constraints provided in the prompt. This setup utilizes the same instruction format established during supervised fine-tuning (detailed in the Methods and Supplementary Information). For this evaluation, we use the same ZINC hard 1k instances described in the previous section. For each instance, we generate and retain up to ten valid and unique SMILES strings per condition. We then compute physicochemical descriptors using the same RDKit-based pipeline applied throughout the benchmark suite. To rigorously quantify success, we define the success rate for each method to reflect the fraction of candidate molecules that simultaneously satisfy every active constraint specified for a given test.

We apply several distinct conditional constraints to test the model’s generative flexibility. *Reasonable* enforces the composite medicinal-chemistry filter used in previous sections, meaning a molecule is only counted if it passes all structural and strain rules, not merely if it is syntactically valid. The *Ro5* condition applies Lipinski-style drug-likeness limits to the fully assembled molecule: a molecular weight of at most 500 Da, a calculated log *P* of at most 5, no more than five hydrogen-bond donors, a maximum of ten hydrogen-bond acceptors, and a topological polar surface area not exceeding 140 Å^2^. To constrain the linker topology itself, we use *Ring* and *Branched* labels, requiring the linker to contain at least one ring or to be classified as branched acyclic, respectively. Furthermore, we apply size limits using *rotB* and *heavy* labels, setting lower bounds that require more than two rotatable bonds and more than six heavy atoms in the linker alone to avoid trivial connections. To isolate the model’s instruction-following capabilities from its unconditional generative capabilities, we compare the prompt-conditioned LinkLlama outputs against an unconditional LinkLlama, as well as the baseline methods DeLinker and DiffLinker, scored with the exact same set of rules for each condition.

Table 2 summarizes the model success rate across various linker and molecular requirements. These results demonstrate LinkLlama’s strong adherence to user prompts, effectively shifting its generative distribution on demand. For the relatively mild Ro5 constraint, the prompted generation performs slightly above the unconditional baseline. This performance is expected, as the unconditional bulk distribution already heavily overlaps with standard drug-like property bands (Supplementary Figure S3.6). However, the performance gap widens significantly when introducing topology constraints. Prompts requiring specific topologies, such as ring or branched constraints, elevate the success rates well above the unconditional background distributions. When evaluating complex, joint constraints in the case of Ring + Ro5 + rotB/heavy + reasonable, LinkLlama retains a prompted success rate in the mid-40% range and meaningfully shift the distribution towards the desired distribution (Supplementary Figure S3.7). In stark contrast, the unconditional LinkLlama and baseline methods collapse to single digits or no success at all under these strict, overlapping criteria. Overall, these results suggest that, under natural language guidance, LinkLlama can reliably navigate complex, multi-objective design spaces that remain inaccessible to unconditional generation or baseline methods.

**Table 2:** Constraint satisfaction rates on the ZINC hard 1k dataset. The table displays the percentage of generated molecules that simultaneously satisfy every active constraint for a given condition. All percentages represent the bootstrap mean ± standard deviation calculated over 2000 bootstrapping samples of the instances.

| Condition | LinkLlama |  | Baselines |  |
| --- | --- | --- | --- | --- |
|  | Conditional | Unconditional | DeLinker | DiffLinker |
| Ro5 + reasonable | 81.2±0.4 | 79.9±0.4 | 21.8±0.4 | 31.8±0.6 |
| Ring + reasonable | 90.4±0.3 | 63.6±0.5 | 0.3±0.1 | 9.5±0.4 |
| Ring + Ro5 + reasonable | 80.1±0.4 | 55.6±0.5 | 0.3±0.1 | 8.2±0.3 |
| Ring + rotB/heavy + reasonable | 49.2±0.6 | 9.0±0.3 | 0.0±0.0 | 1.1±0.1 |
| Ring + Ro5 + rotB/heavy + reasonable | 43.1±0.5 | 7.4±0.3 | 0.0±0.0 | 0.9±0.1 |
| Branched + reasonable | 79.9±0.4 | 22.5±0.4 | 2.1±0.2 | 24.5±0.5 |
| Branched + Ro5 + reasonable | 76.7±0.5 | 21.6±0.4 | 2.1±0.2 | 23.2±0.5 |
| Branched + Ro5 + rotB/heavy + reasonable | 68.1±0.5 | 15.8±0.4 | 0.0±0.0 | 6.2±0.3 |
| Branched + rotB/heavy + reasonable | 70.6±0.5 | 16.6±0.4 | 0.0±0.0 | 6.7±0.3 |

### Benchmarking on 3D-Based HiQBind Dataset

To further validate our model on a more challenging and structurally relevant benchmark, we utilize the HiQBind dataset^36^, which consists of high-quality PDB chains paired with experimental crystal ligand poses. Because the structural fragments in this dataset form desirable interactions with target proteins, they serve as highly realistic starting points for fragment-based drug design. We evaluate the models on 1k random and 1k hard complexes, with the hard split enforcing the identical linker-difficulty rules defined previously (i.e., requiring linkers to contain more than three rotatable bonds or more than seven heavy atoms). It is important to note that we employ unconditional LinkLlama generation in this section to establish a rigorous baseline comparison. In practice, a user with a specific structural hypothesis could leverage our prompt-conditioning capabilities to combine various design properties, explicitly tailoring the generated linkers to the unique chemistry of the pocket.

As detailed in Supplementary Table 2 and Figure 4, we observe generative performance and 3D structural trends consistent with our earlier ZINC benchmarking. Although overall QED scores shifts slightly lower across all methods on this harder dataset, LinkLlama maintains near-perfect validity and high uniqueness and demonstrates a stark advantage in structural reasonability, reaching 80.9% on the hard split compared to DiffLinker’s mid-20% mean pass rate. This divergence in chemical realism is heavily reflected in the 3D geometry evaluations. Again, while DiffLinker achieves competitive pose agreement (mean RMSD), it does so at a severe energetic cost, exhibiting a heavy-tailed distribution of high internal strain (Δ*E*). Conversely, LinkLlama balances competitive RMSD with relaxed, low-strain geometries, confirming its ability to reliably produce physically viable and chemically reasonable ligand structures even within complex, protein-bound environments.

**Figure 4:**
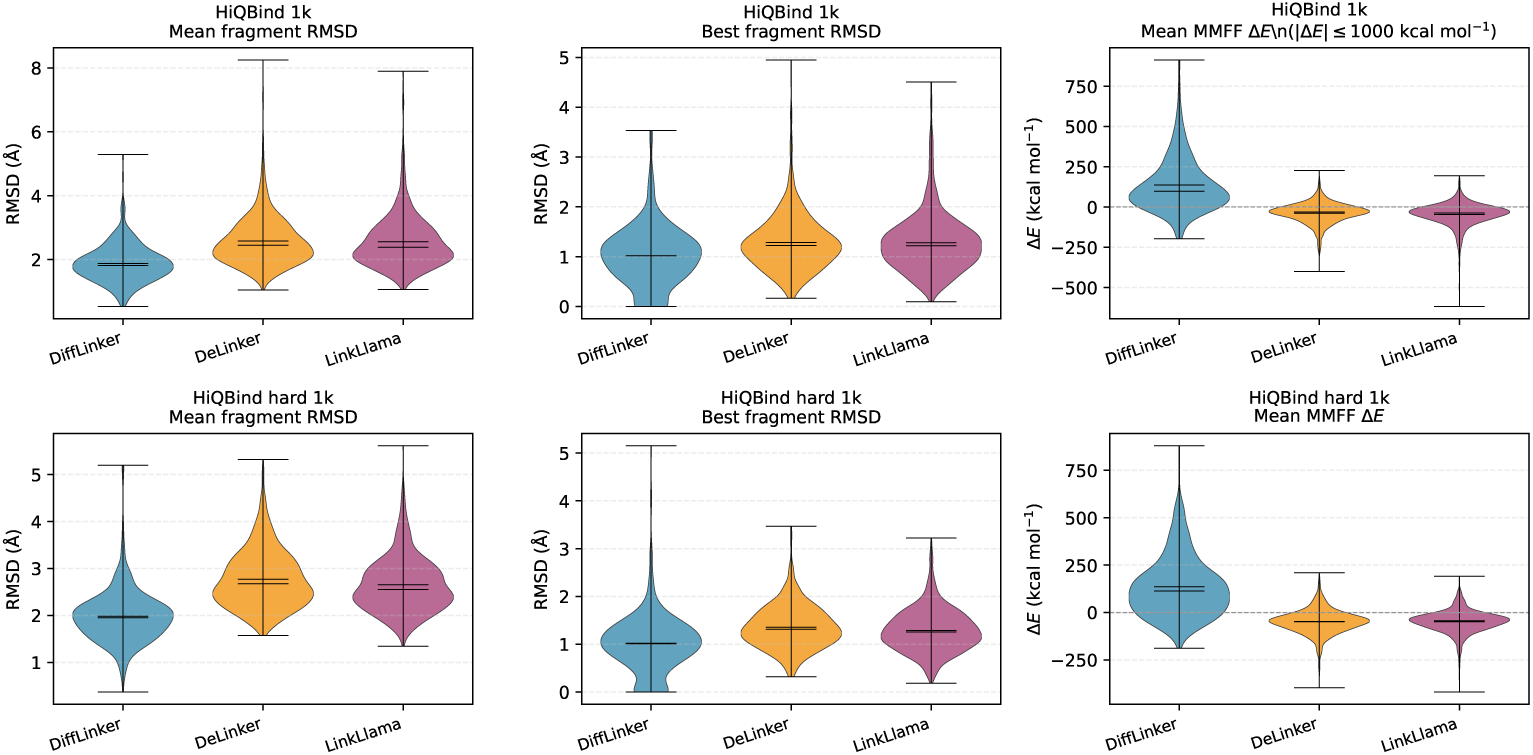
Assessment of 3D geometric realism and internal strain on the HiQBind dataset. The violin plots display per-instance distributions for mean fragment RMSD, best fragment RMSD, and mean MMFF Δ*E* (kcal/mol), following the layout of Figure 3. For the HiQBind random 1k plot specifically, the Δ*E* column filters out extreme outliers, retaining only instances with |Δ*E*| ≤ 1000 kcal/mol as an MMFF diagnostic cutoff. Both the HiQBind hard plots and all RMSD panels incorporate every valid generated instance.

To evaluate functional binding potential, we use Uni-Dock^37^ to dock valid proposed ligands into their respective protein pockets and calculate the per-instance mean docking score differences (Δ) relative to the experimental reference pose. Detailed procedures are include in Supplementary Information S1. As shown in Figure 5, although most of the poses in all methods do not outperform the reference ligand, there still are some portions of all model’s distribution falls into the negative delta range. This observation indicates that the model routinely proposes novel, strong-binding linker conformations that match or improve upon the binding affinities of the native ligands within the actual pocket environment, suggesting that it can not only link the two fragments under such context, but also links the fragments in a physically meaningful way for generating good whole-molecule binders.

**Figure 5:**
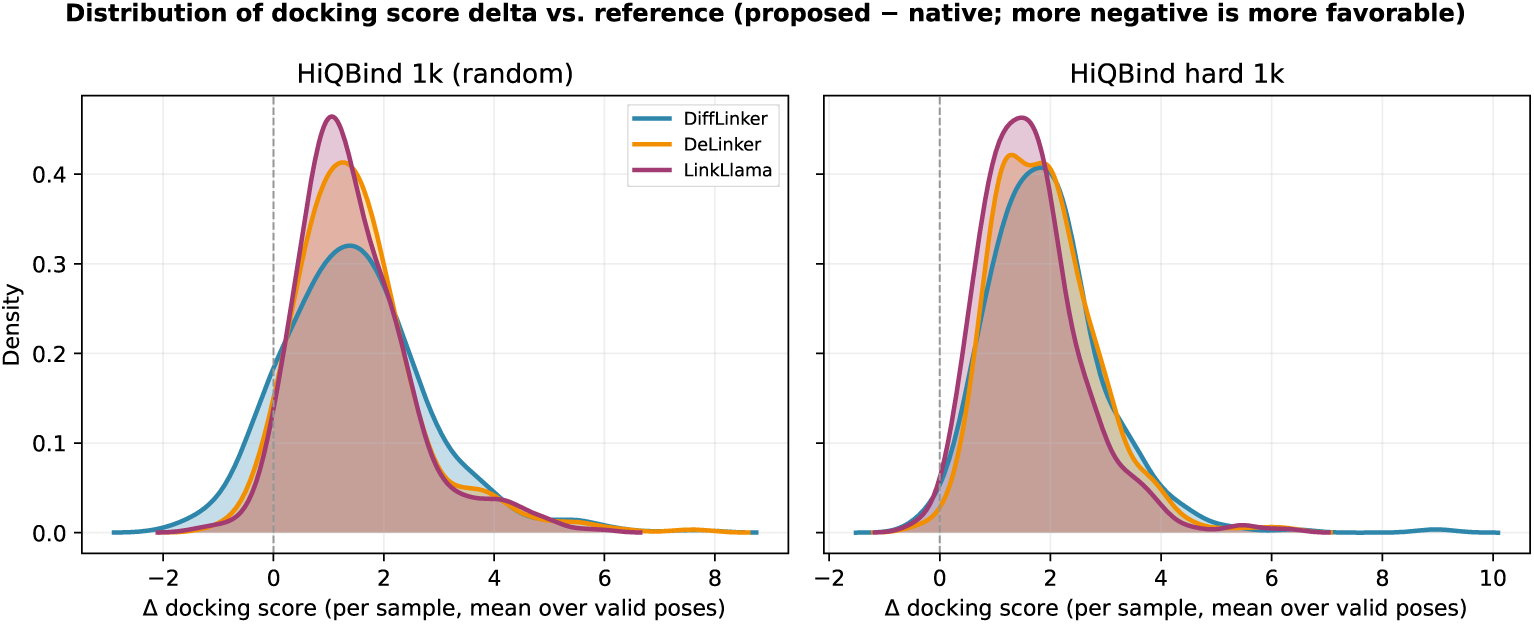
Distribution of mean docking score differences relative to the reference ligands. The plot displays kernel density estimates of the per-sample mean docking score delta (calculated as the proposed score minus the reference score) on valid instances for the HiQBind random and hard 1k splits.

### Case Studies: Scaffold Hopping and PROTAC Design

To demonstrate the practical utility of *LinkLlama* in a realistic drug discovery scenario, we investigate a specific case from the HiQBind dataset of the mineralocorticoid receptor (MR) complexed with a potent antagonist (PDB ID: 6L88)^38^. The reference ligand exhibits a distinct binding mode that its terminal sulfonyl moiety forms a crucial hydrogen bond with Arg817, accessing a previously unexplored sub-pocket in the crystal structure. Simultaneously, its trifluoromethylbenzyl side chain occupies an adjacent small hydrophobic cavity. Recognizing an opportunity for scaffold hopping, we sought to replace the central core to discover novel chemical matter that maintains these critical anchoring interactions.

Although the reference molecule is highly optimized with a favorable docking score (−9.7 kcal/mol), we use LinkLlama to unconditionally sample 100 diverse linker candidates. These generated molecules are then rigorously filtered so that they have improved docking scores relative to the reference, low fragment RMSD (*≤* 2Å) to preserve the required 3D vectors of the terminal groups, and high chemical reasonability to pass all five checks. To rigorously evaluate binding stability, the top 10 designs are then subjected to 200 ns molecular dynamics (MD) simulations.

As shown in Figure 6, LinkLlama successfully generalizes to this complex task. As shown in Figure 6, The model proposed novel, synthetically accessible heterocyclic cores, such as substituted isoxazoles and diazoles, that yields more favorable docking scores than the reference. Structural overlay confirms that the generated scaffolds perfectly maintain the trajectory of the functional terminal groups. Furthermore, the MD trajectories reveal that the generated ligands remain highly stable over the 200 ns duration, exhibiting mean ligand RMSDs (1.30–1.65 Å) that are tighter than or comparable to the reference ligand (1.70 Å). This observation confirms that LinkLlama can reliably produce stable, synthetically realistic candidates that improve upon known potent inhibitors.

**Figure 6:**
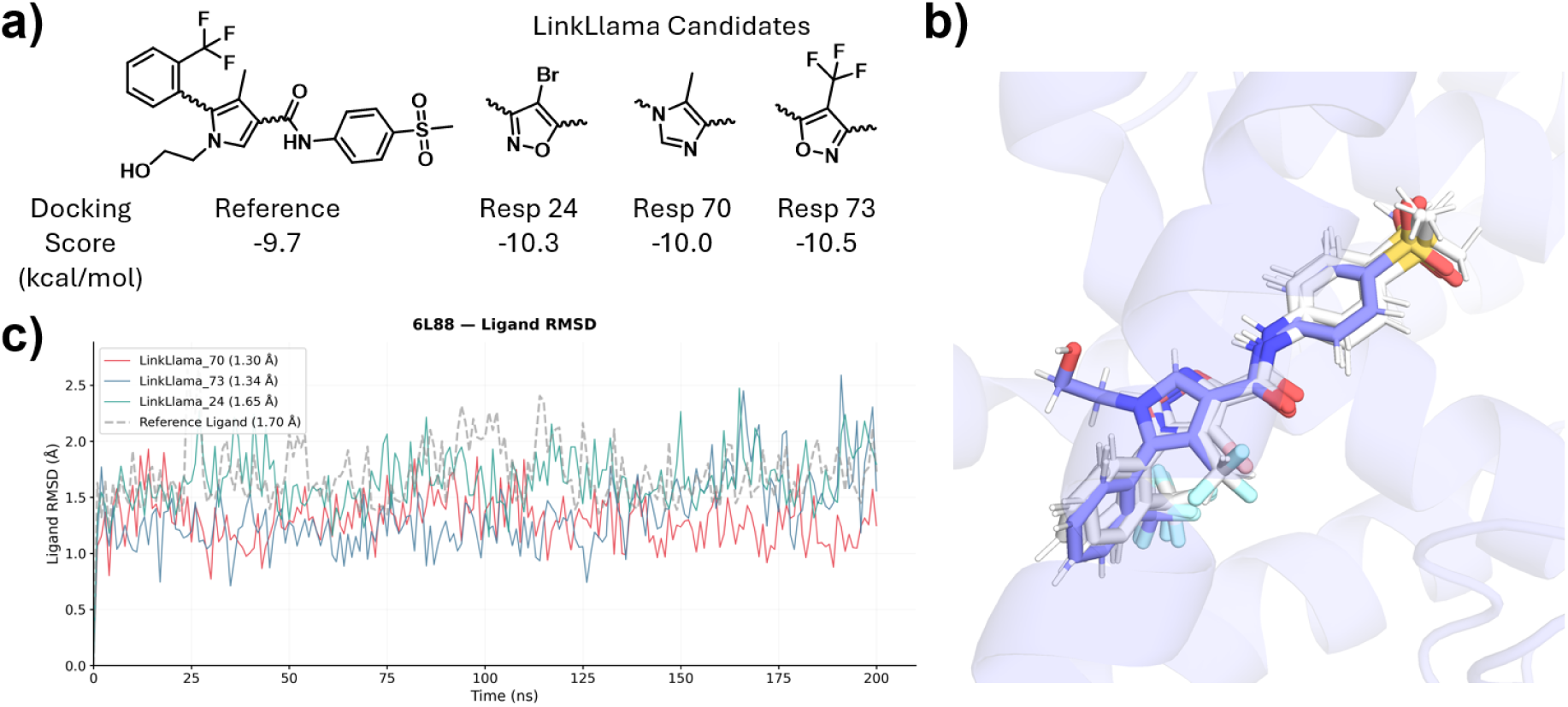
Scaffold hopping case study on the mineralocorticoid receptor (PDB ID 6L88). (a). 2D chemical structures of the reference ligand and three top-performing LinkLlama candidates (Resp 24, 70, and 73), illustrating the successful generation of diverse heterocyclic cores that improve upon the reference docking score. (b). 3D structural overlay of the docked poses of LinkLlama candidates on top of crystal ligand pose.(c). Ligand root-mean-square deviation (RMSD) over 200 ns molecular dynamics simulations.

We also demonstrate LinkLlama’s utility in proteolysis-targeting chimera (PROTAC) linker design. PROTACs are heterobifunctional molecules consisting of two ligands joined by a linker: one recruits a protein of interest (POI) and the other recruits an E3 ubiquitin ligase^39^. Unlike standard inhibitors, where efficacy is primarily driven by ligand binding affinity, PROTAC activity relies heavily on the linker’s ability to stabilize a cooperative ternary complex (POI–PROTAC–Ligase). Consequently, once the warhead and E3 anchoring ligands are identified, designing a linker with the optimal length, composition, and geometry becomes the critical bottleneck^39^. For this case study, we selected a well-characterized BRD4–VHL PROTAC from PROTAC-DB 3.0^40^ (PDB ID: 6SIS^41^).

In the original 6SIS structure, the PROTAC utilizes a complex macrocyclic architecture. As shown in Figure 7(a), the VHL E3 ligand is shown in blue, and the BRD4 POI ligand is shown in green. We therefore task LinkLlama with generating linear linker alternatives to replace the specific bridging segment highlighted in red, aiming to simplify synthetic complexity while maintaining the vital ternary interaction. Given the expansive chemical space of PROTACs, we generate approximately 300 candidates using diverse property-constraining prompts to prioritize chemical reasonability (Supplementary Information S1). Valid designs are docked into the ternary interface, and we select molecules that exhibited low fragment RMSDs compared to the experimental crystal poses of the anchoring ligands. The top candidates are then evaluated using 200 ns MD simulations.

**Figure 7:**
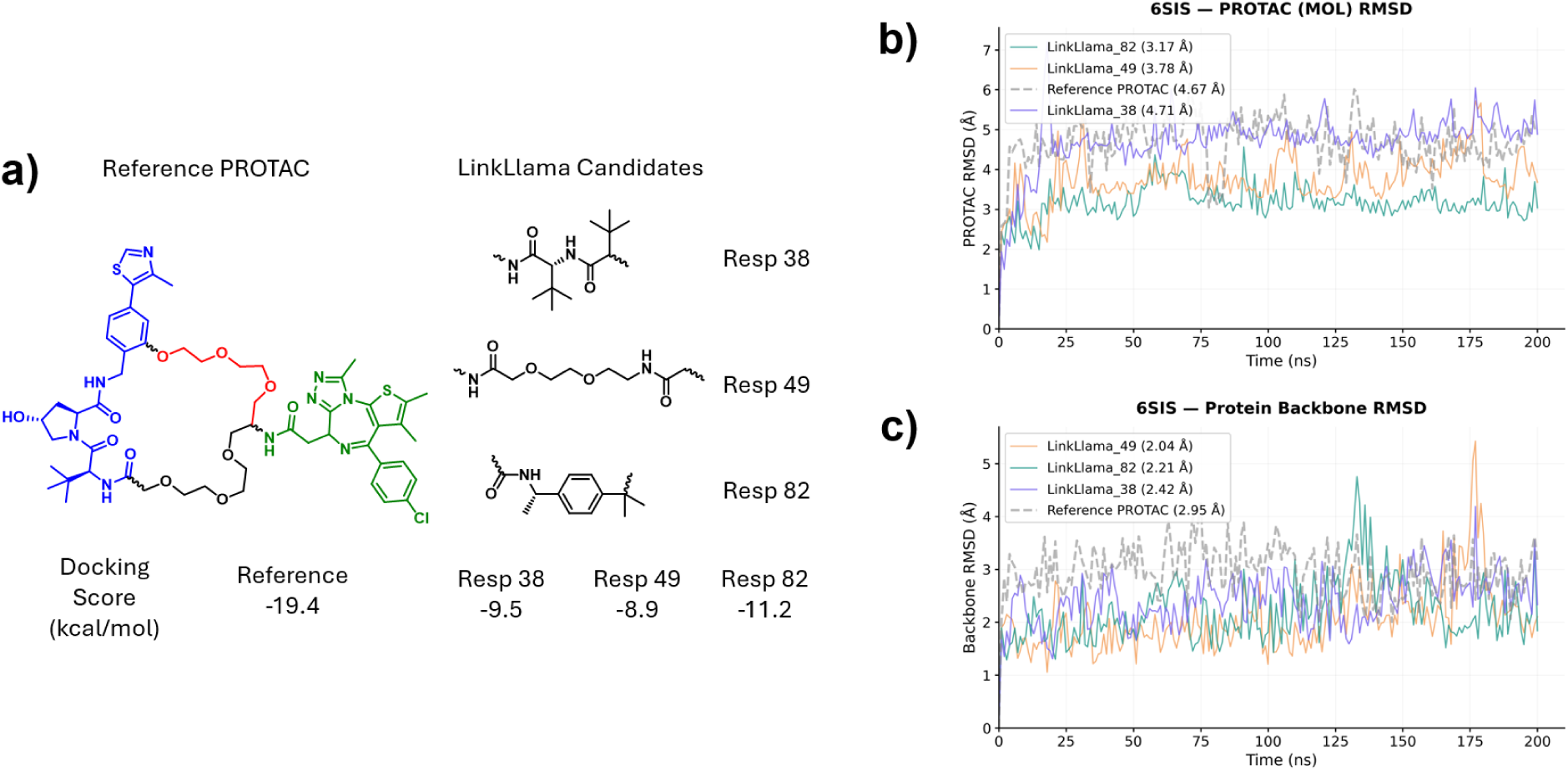
PROTAC linker design for the BRD4-VHL ternary complex (PDB ID 6SIS). (a). 2D chemical structures of the reference macrocyclic PROTAC (targeted linker region in red, VHL ligand in blue, and BRD4 ligand in green) alongside three linear LinkLlama candidates (Resp 38, 49, and 82) and their corresponding docking scores. (b). PROTAC ligand RMSD over 200 ns MD simulations. (c). Protein backbone RMSD over the simulation.

As shown in Figure 7(a), LinkLlama successfully generates diverse linear linkers, incorporating peptidic, PEG-based, and rigid aryl motifs. While these designs bypass the synthetic burden of macrocyclization, MD trajectories demonstrate excellent conformational stability. The generated PROTACs exhibit ligand RMSDs (3.17–4.71 Å) that are comparable to or significantly tighter than the reference macrocycle (4.67°A). Crucially, the overall ternary complex stability is enhanced by these new linkers as the protein backbone RMSD for the generated candidates (2.04—2.42 Å) outperforms the reference complex (2.95 Å) throughout the trajectory. These results underscore the model’s capacity to navigate complex, long-range spatial constraints and propose viable PROTAC candidates. Overall, both cases demonstrate LinkLlama’s utility for linker proposal with various fragment starting points.

## Discussion

In this study, we introduced LinkLlama, an LLM-based framework that reframes the fragment-linking problem as conditional SMILES string generation task. By leveraging supervised finetuning, LinkLlama successfully turns a general-purpose LLM into an expert model capable of navigating the accessible, drug-like chemical space that previous models often struggle to capture. By aligning the chemical reasonability in its training input prompts with the generated linkers, LinkLlama acts as a rational “slicer” of chemical space, successfully producing novel yet diverse linkers for various fragment inputs while strictly maintaining their chemical rationality. Our findings indicate a significant leap in chemical quality, achieving 70–80% chemically reasonable linker designs across various datasets, effectively doubling the performance of previous baseline methods. This capability suggests that medicinal chemists are far more likely to work with the raw hypotheses proposed by LinkLlama due to their superior chemical fidelity.

Another defining advantage of LinkLlama is its capacity for reinforcement learning-free conditional sampling via direct prompt manipulation. Generative models have historically relied on computationally expensive and often unstable RL cycles to achieve multi-objective optimization^11,25,26^. LinkLlama replaces this paradigm with an “alignment-by-design” approach. By steering the model through natural language prompts, medicinal chemists can seamlessly tailor chemical property distributions without engineering complex reward functions. This setup lowers the computational barrier for generating actionable leads while maintaining the spatial integrity of fragment hits across diverse modalities. Whether the task involves simple fragment joining, scaffold hopping, or navigating the demanding architectural constraints of PROTAC design, LinkLlama consistently outputs chemically reasonable designs that demonstrate strong computational validation.

Looking forward, the utility of frameworks like LinkLlama is further underscored by the rapidly shifting landscape of structural biology. With continuous advancements in high-throughput Xray crystallography and cryo-EM like the OpenBind Initiative^42^, experimental fragment hit data is becoming increasingly abundant. This wealth of structural data is driving a paradigm shift away from purely *de novo* whole-molecule generation toward a more grounded, iterative pipeline: fragment identification, experimental validation, generative linking, and subsequent validation. In line with the recent ASAP-Polaris-OpenADMET challenge ^43^ demonstrating that fragment anchors can effectively drive accurate pose predictions^44^, LinkLlama capitalizes on these high-confidence spatial constraints. By reliably bridging experimentally validated fragments, it maximizes the value of structural data and ensures that generative design remains tethered to experimental reality.

In another aspect, LinkLlama is designed to function not in isolation, but as a foundational generative engine within a broader LLM-based chemical design ecosystem. Conceived as part of the “Chemical Llama Suite,” it works synergistically with models like SmileyLlama for general molecular generation and SynLlama for retrosynthetic planning. Because these models communicate natively via standard SMILES and natural language, they are perfectly positioned for integration with autonomous AI agents^18,19,45^. We envision a “closed-loop” discovery workflow where an agentic system autonomously identifies promising candidates via LinkLlama, evaluates them through automated docking, FEP, or ADMET simulations, and continuously refines the model’s output using techniques such as Direct Preference Optimization (DPO). This architecture effectively achieves an LLM-based equivalent of the REINVENT tool suite^46^, transforming the generative model from a static tool into a dynamic, evolving partner in the transition from isolated fragments to high-quality clinical leads.

## Supplementary Information

Details of ChEMBL36 dataset preparation, molecular fragmentation procedure, SFT corpus preparation, docking and MD procedures, and additional analysis are provided in the Supplementary Information.

## Data and Code Availability

All the codes are available on GitHub: https://github.com/THGLab/LinkLlama

## Author Contributions

KS and THG conceptualized and defined the goal of the project. KS wrote the code and trained the models. KS, YW, and JP performed analysis for the Results section. KS and THG wrote the paper. All authors discussed the results and made comments and edits to the manuscript.

## Acknowledgments

This work was supported by National Institute of Allergy and Infectious Disease grant U19AI171954. This research used computational resources of the National Energy Research Scientific Computing, a DOE Office of Science User Facility supported by the Office of Science of the U.S. Department of Energy under Contract No. DE-AC02-05CH11231.

## Supplementary Information

### S1 Additional Methodology Details

#### Dataset Curation and Fragmentation Protocol

To ensure the quality and drug-like relevance of the parent pool prior to fragmentation, the initial dataset of 2,854,815 raw molecules sourced from ChEMBL36 undergoes a rigorous, multi-step filtering cascade. Molecules are systematically excluded based on standard physicochemical boundaries and structural viability using RDKit^28^. The sequential filtering protocol results in the removal of compounds based on the following criteria:

- **Validation and Salt Removal:** Molecules failing initial graph construction, failing salt-stripping protocols, or failing standard sanitization checks are removed (1 molecule).
- **Atom Type Restrictions:** Compounds containing non-standard elemental species or non-physiological charge states are excluded. Allowed atom types are strictly limited to the following symbol, valence, and formal charge combinations: C4(0), O2(0), N3(0), F1(0), S2(0), Cl1(0), S6(0), N4(+1), O1(−1), Br1(0), P5(0), and I1(0) (37,213 molecules removed).
- **Minimum Size Constraint:** Extremely small fragments lacking sufficient complexity for linker derivation, defined as having fewer than 6 heavy atoms, are removed (587 molecules).
- **Maximum Molecular Weight:** Exceptionally large macromolecules or polymers with a molecular weight of *≥* 1500 Da are discarded (23,957 molecules).
- **Ring System Constraints:** Molecules with excessively complex topologies, specifically those containing more than 10 total rings or more than 8 aromatic rings, are excluded (4,505 molecules removed).
- **Carbon Ratio:** Compounds with an unusually low fraction of carbon atoms (defined as a carbon-to-heavy-atom ratio of *≤* 0.5), which often indicate highly inorganic or non-drug-like frameworks, are removed (25,659 molecules).

Following this filtering cascade, the surviving structures are canonicalized and deduplicated. This yields a final robust pool of 2,665,082 unique, valid parent molecules (representing 93.4% of the initial input), which subsequently serve as the foundation for the fragmentation and supervised fine-tuning (SFT) corpus.

#### Computational Fragmentation and Structural Filtration

The generation of training triplets (fragment-linker-fragment) is executed using the Matched Molecular Pair Analysis (MMPA) fragmentation algorithm in RDKit. The procedure is designed to isolate realistic, synthetically accessible linker units by strictly enforcing a two-cut mechanism per parent molecule.

Bonds to cleave are defined by the SMARTS^32^ [#6+0;!$(*=,#[!#6])]!@!=!#[*]. This specific query restricts fragmentation to acyclic single bonds attached to a neutral carbon atom. The pattern utilizes recursive SMARTS exclusions to prevent the disruption of complex substitution environments or conjugated systems.

Following MMPA decomposition, the resulting structural triplets are parsed into a central core (the linker) and terminal chains (the fragments). These isolated components are subjected to a rigorous filtration cascade. A fragmentation event is considered successful and added to the dataset only if all the following conditions are simultaneously satisfied:

- Minimum Size Requirements: The central linker unit must contain a minimum of 3 heavy atoms, and each separate terminal fragment must contain a minimum of 5 heavy atoms.
- Attachment Point Validation: The isolated linker subgraph must possess exactly two dummy atoms (atomic number equal to 0), confirming it represents a valid two-point connection bridge rather than a terminal appendage.
- Topological Separation: The shortest path between the two dummy atoms on the linker subgraph is calculated to ensure adequate spatial separation between the fragments. We calculate the topological distance as the total number of bonds in the shortest path minus two (accounting for the dummy bonds). The system requires a minimum internal path length of 2.
- Mass Balance: To prevent the generation of skewed decompositions where the linker dominates the molecular framework, the number of heavy atoms in the linker must be less than or equal to the heavy atom count of the smallest terminal fragment.

Molecules that fail to produce a valid double-cut matching the specified SMARTS pattern naturally fall back to an empty decomposition state and are subsequently removed from the training corpus, along with any decompositions that fail the topological or size-based filters.

#### The Chemical Reasonability Workflow

As mentioned in Main, a fragmentation case is reasonable if and only if all 5 chemical reasonability checks pass. We detail the five evaluation criteria below.

- Bridgehead ring structure (Linker). Fail if any linker heavy atom belongs to three or more rings in the smallest-set-of-rings decomposition, flagging a single center shared by too many ring systems.
- Uncommon ring systems (Linker). Fail if any ring system in the linker has an occurrence count below 100 in the full ChEMBL36 parent database used to build the training pool.
- Undesirable SMARTS (Molecule). Fail the full molecule matches to the curated list of undesirable SMARTS, including cyclopentadiene, cyclopentadiene ylidenes, aromaticity-breaking tautomers, antiaromatic system, unstable halogen-heteroatom bonds, unstable fused rings, allenic system, thiazyl linkages, and peroxide bonds.

**–** [C^2]1=[C^2]-[C^2]=[C^2]∼[C;!d4]∼[C;!^2;d2]1

**–** [C^2]1∼[C^2]∼[C^2]∼[C^2]∼[C;!^2;d2]∼[N]1

**–** [#6^2]1∼[#6^2]∼[#6^3;!d4]∼[#6^2]2∼[#6^2]∼

[#6^2]∼[#6^2]∼[#6^2](∼[*])∼[#6^2]∼2∼[#6^2]∼1

**–** [#6]1(=[*])[#6]=[#6][#6]=[#6]1

**–** [#6]1=[#6][R{2-}]=[R{2-}]1

**–** [#6^2]1∼[#6^2]∼[#6^2]∼[#6^2]∼[#6^1]∼[#6^1]∼1

**–** [#7,#8,#16]-[#9,#17,#35,#53]

**–** [r3,r4]@[r5,r6]

**–** [*]=[#6,#7,#8]=[*]

**–** [#7,#16]=[#16]

**–** [#8]-[#8]

In addition to the patterns mentioned above, this check will fail if the generated pyrroles are not in one of the following correct forms.

**–** [N^2]1∼[C,N;^2](=[*])∼[C,N;^2]∼[C,N;^2]∼[C^3]1

**–** [N^2]1∼[C,N;^2]∼[C,N;^2](=[*])∼[C,N;^2]∼[C;^3]1

**–** [N^2]1∼[C,N;^2]∼[C,N;^2]∼[C,N;^2](=[*])∼[C;^3]1

**–** [C,N;^2](=[*])1∼[N;^2]∼[C,N;^2]∼[C,N;^2]∼[C;^3]1

**–** [C,N;^2]1∼[N;^2]∼[C,N;^2](=[*])∼[C,N;^2]∼[C;^3]1

**–** [C,N;^2]1∼[N;^2]∼[C,N;^2]∼[C,N;^2](=[*])∼[C;^3]1

- PAINS (Molecule). Fail if the full molecule matches to the standard PAINS structural-alert catalog.
- Brenk filters (Molecule). Fail if the full molecule matches to the Brenk medicinal-chemistry rule set.

#### Preparation of LLM Fine-Tuning Corpus

For all the training data of selection, we apply the same instruction prompt as follows.

You are an expert medicinal chemist specializing in linker design. Your task is to design a linker to connect given fragments and deduce whether the final molecule is chemically reasonable. The output should be in JSON format.

For each input-output training pairs, after collecting their fragment information, linker and molecular properties, and the reasonability checks, we structure these data into the following formats.

FRAGMENT INFO Given the above information about the fragments and attachment points, design a LINKER TYPE linker LINKER PROPERTIES to connect them. The final molecule should be REASONABILITY.MOLECULE PROPERTIES

The placeholders are substituted during training. FRAGMENT INFO field is replaced with the fragment SMILES strings, and distance and angle between the two fragments. The REASONABILITY field is always either reasonable or unreasonable according to the overall reasonability check. All the other field in the input has a 50% chance of showing up in the training script to ensure our training robustness. The specific allowed input for these additional field is shown in Supplementary Table S2.1.

The corresponding output line is a JSON object with keys linker (SMILES of the training linker) and reasoning (human-readable pass/fail summary for the five reasonability checks) as illustrated in Figure 1.

#### Supervised Fine-Tuning

After preparing the aforementioned training data into the JSONL format, we fine-tune Llama-3.2-1B-Instruct model (1 Billion parameters) ^47^ using the Axolotl package ^48^ for 1 epoch. We apply Low-Rank Adaptation (LoRA) with a rank of *r* = 32 and *α* = 16 to the linear layers of the model ^49^. We use FlashAttention-2^50^, with the Adam optimizer^51^, crossentropy loss, and a cosine learning rate scheduler with a maximum learning rate of 2 *×* 10*^−^*^4^. The training time for our final selected model requires four A100 GPUs on one node for approximately 6.5 hours.

#### HiQBind/PROTAC docking procedures using Uni-Dock

For the HiQBind and PROTAC docking study, ligands were taken from the model-generated ensembles after the same validity screen used elsewhere in our linker benchmarks. For each protein target, we use the paired refined receptor PDB and co-crystallized reference ligand SDF as the center of the docking box. The box center is the midpoint of the ligand heavy-atom coordinate ranges, and each axis length is the larger of 24 Å and the ligand bounding span plus a 5 Å padding margin. The receptor is cleaned to coordinate records, converted to PDBQT with Meeko ^52^ using that same center and size, and treated as rigid during docking. Each accepted SMILES is embedded in 3D, prepared as PDBQT with Meeko, and docked with Uni-Dock in fast search mode against the rigid receptor within the defined box. The reported pose corresponds to the top-ranked output, with the docking score read from the PDBQT REMARK fields.

#### Molecular dynamics simulations via GROMACS^53^

The MD protocol used here follows the iMiner protocol ^29^. It initializes with the top-scoring docked pose for each molecule, which is solvated and neutralized with Na^+^ or Cl*^−^* ions. The system is parameterized using the AMBER14SB^54^, GAFF2^55^, and TIP3P force fields for the protein, ligand, and water, respectively. Following energy minimization, the complex undergoes successive 1-ns equilibrations in the NVT and NPT ensembles at 298.15 K and 1 bar. Then the final production is run for 200 ns for a full simulation and downstream analysis.

#### Prompts for generating PROTAC linker candidates

To sample diverse PROTAC linkers, we apply the following six prompt combinations and generate 50 samples for each prompt.

- **Unconditional:** reasonable.
- **Large MW / HBA:** reasonable; hbd *≤* 5; hba *>* 10; mw *>* 700.
- **Lipophilic–polar:** reasonable; hbd *≤* 5; hba *>* 10; mw *>* 700; log *P >* 6; TPSA *>* 200.
- **Ring linker:** reasonable; ring-containing; hbd *≤* 5; hba *>* 10; mw *>* 700.
- **Branched linker:** reasonable; branched; hbd *≤* 5; hba *>* 10; mw *>* 700.
- **Chain flexible:** reasonable; chain; rotb *≥* 4; heavy atoms *≥* 7; hbd *≤* 5; hba *>* 10; mw *>* 700.

## S2 Supplementary Tables

**Table S2.1:**
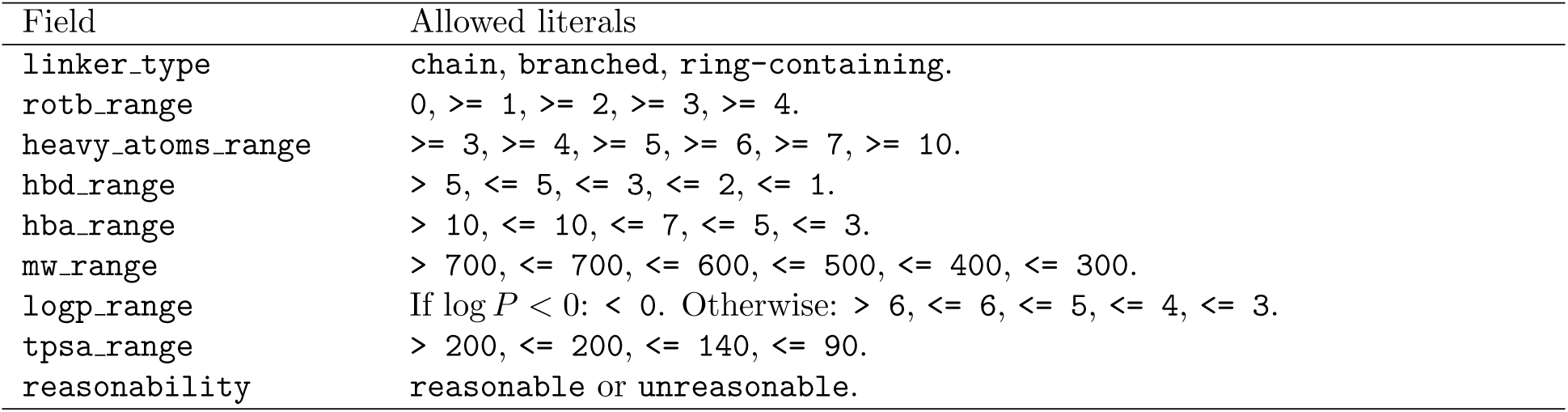
Configurable prompt fields and allowed input strings for linker and molecule property clauses. LogP uses the log *P <* 0 branch only when the molecule’s computed log *P* is negative.

**Table S2.2:** Generative performance on the HiQBind random 1k and hard 1k benchmarks. Conventions match Table 1: validity, uniqueness, novelty, and reasonability are reported as percentages, while QED and SA are dimensionless. Percentage columns display the instance-aggregated mean ± bootstrap standard deviation of that mean. QED and SA represent the instance-mean ± standard deviation across instances. Novelty values for the baseline methods are excluded due to differing training set derivations.

| Split | Model | Validity | Uniqueness | Novelty | QED | SA | Reasonability |
| --- | --- | --- | --- | --- | --- | --- | --- |
| Random 1k | DeLinker | 94.6±0.4% | 87.4±0.2% | — | 0.44±0.18 | 3.84±0.88 | 35.6±1.1% |
| Random 1k | DiffLinker | 87.2±1.2% | 64.2±1.2% | — | 0.53±0.16 | 3.73±0.98 | 18.3±1.0% |
| Random 1k | LinkLlama | 99.8±0.0% | 83.9±0.4% | 32.4±0.5% | 0.43±0.20 | 3.83±0.93 | 53.4±1.5% |
| Hard 1k | DeLinker | 92.6±0.6% | 88.2±0.2% | — | 0.44±0.16 | 3.73±0.75 | 44.0±0.9% |
| Hard 1k | DiffLinker | 88.5±1.1% | 60.9±1.1% | — | 0.52±0.14 | 3.56±0.87 | 25.7±1.1% |
| Hard 1k | LinkLlama | 99.8±0.0% | 84.7±0.5% | 40.1±0.5% | 0.44±0.17 | 3.62±0.75 | 80.9±0.9% |

## S3 Supplementary Figures

**Figure S3.1:**
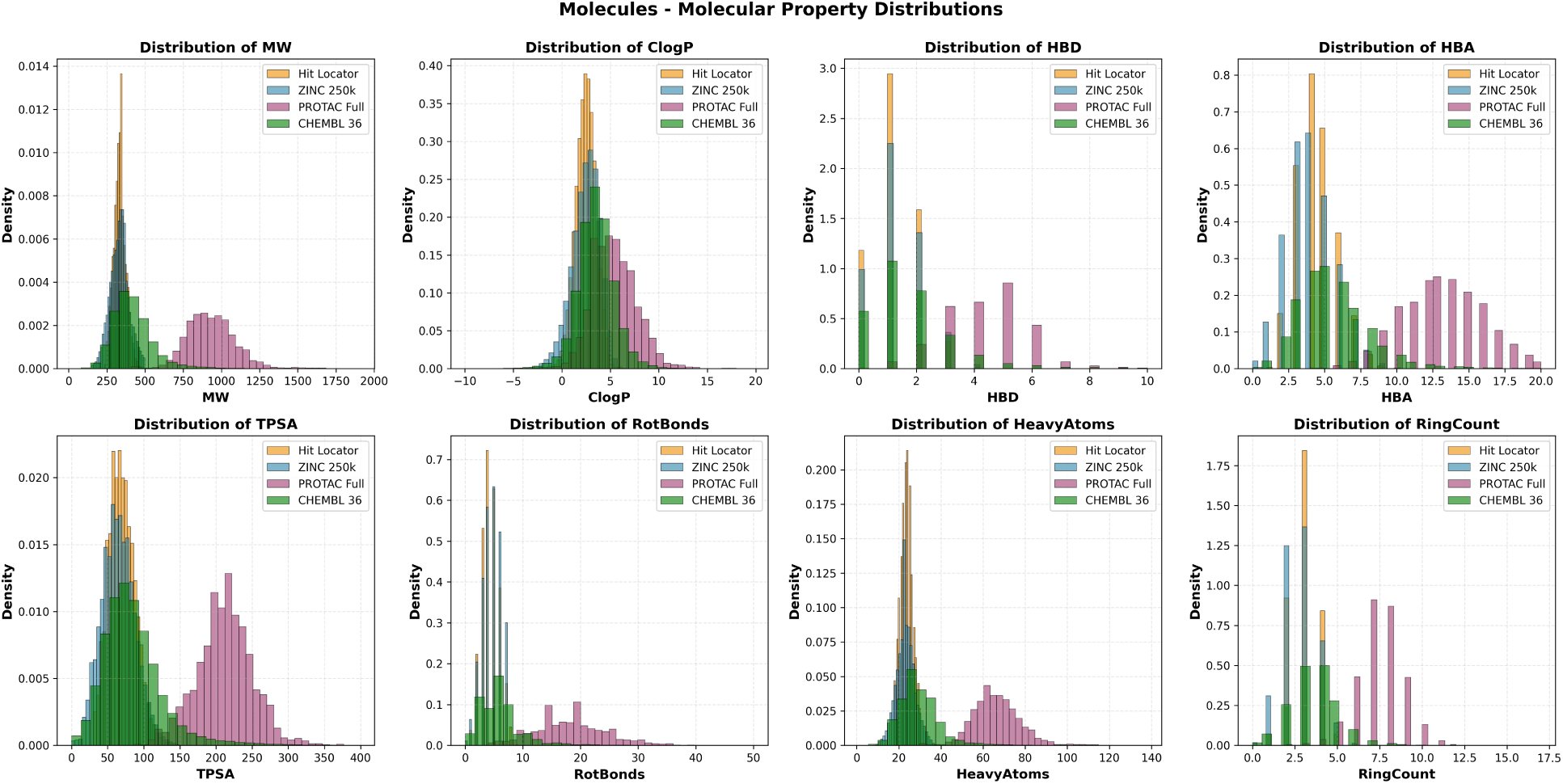
Molecular property distributions comparing the ChEMBL 36 dataset against ZINC 250k, Hit Locator, and a curated PROTAC dataset. Evaluated properties include Molecular Weight (MW), ClogP, Hydrogen Bond Donors (HBD), Hydrogen Bond Acceptors (HBA), Topological Polar Surface Area (TPSA), Rotatable Bonds (RotBonds), Heavy Atoms, and Ring Count. ChEMBL 36 exhibits a significantly broader coverage of chemical space, effectively bridging the gap between traditional small-molecule libraries (ZINC, Hit Locator) and larger chimeric molecules (PROTACs). This comprehensive diversity justifies its selection as the parent pool for LinkLlama’s training corpus.

**Figure S3.2.**
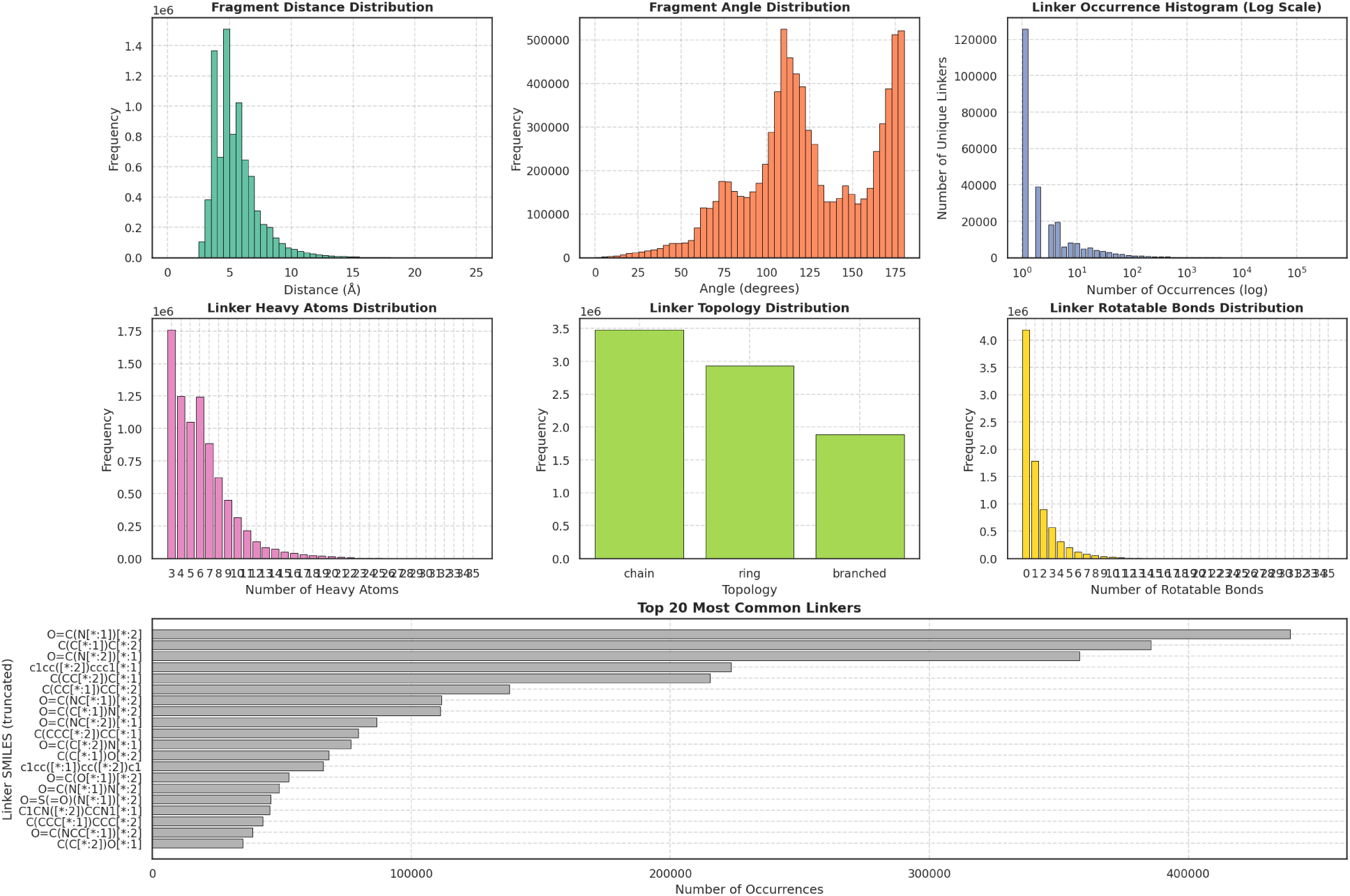
Fragment and linker property distributions for the full ChEMBL36 fragmentation pool. The top right panel (Linker Occurrence Histogram) and bottom panel (Top 20 Most Common Linkers) illustrate a severe long-tail distribution, where a small fraction of simple, highly prevalent linkers dominates the dataset. This extreme imbalance motivates the need for frequency-capping strategies during the curation of the supervised finetuning corpus to prevent model collapse toward common structural motifs.

**Figure S3.3:**
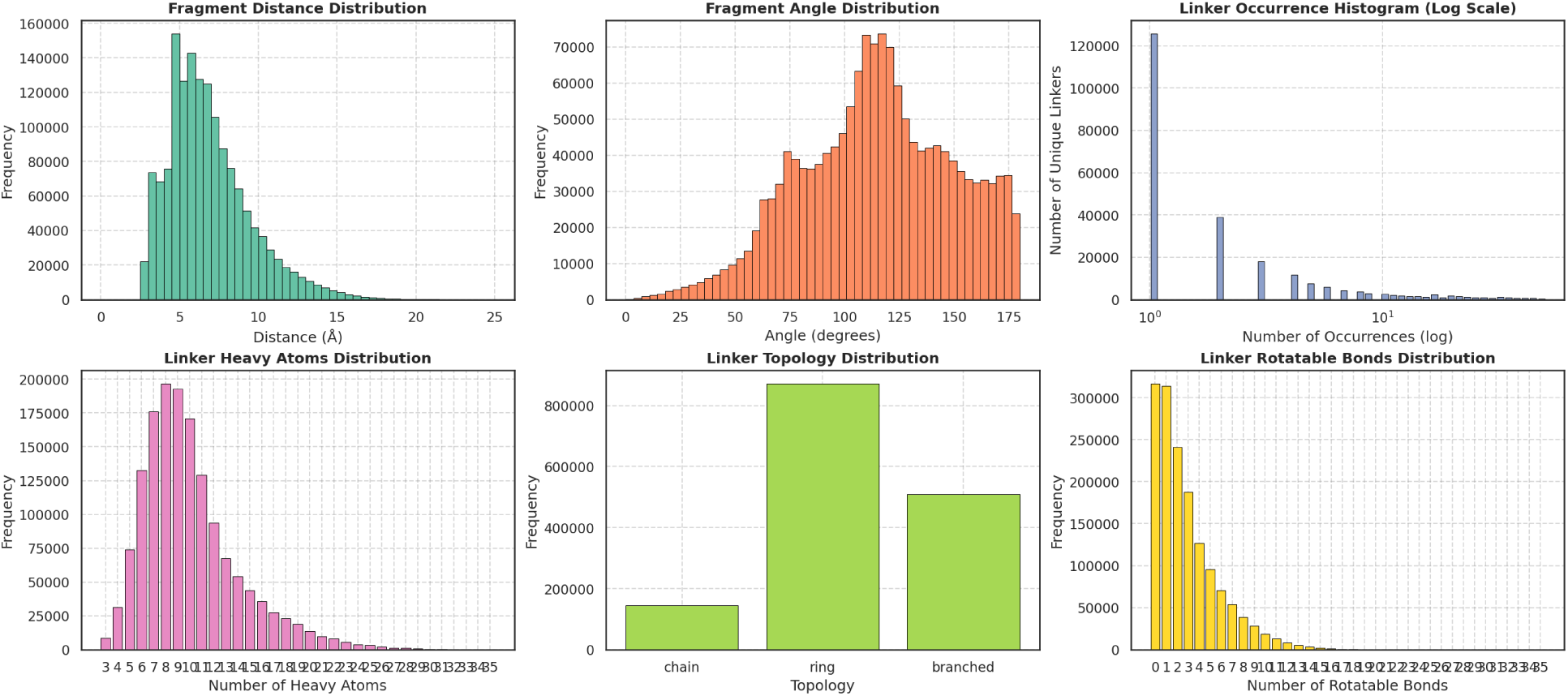
Fragment and linker property distributions for the strictly balanced dataset (Cap50), where the maximum occurrence of any unique linker is restricted to a threshold of 50 examples. While this approach effectively flattens the occurrence histogram and maximizes linker diversity, it slightly shifts the distributions of heavy atoms and rotatable bonds toward more complex topologies compared to the full parent pool.

**Figure S3.4:**
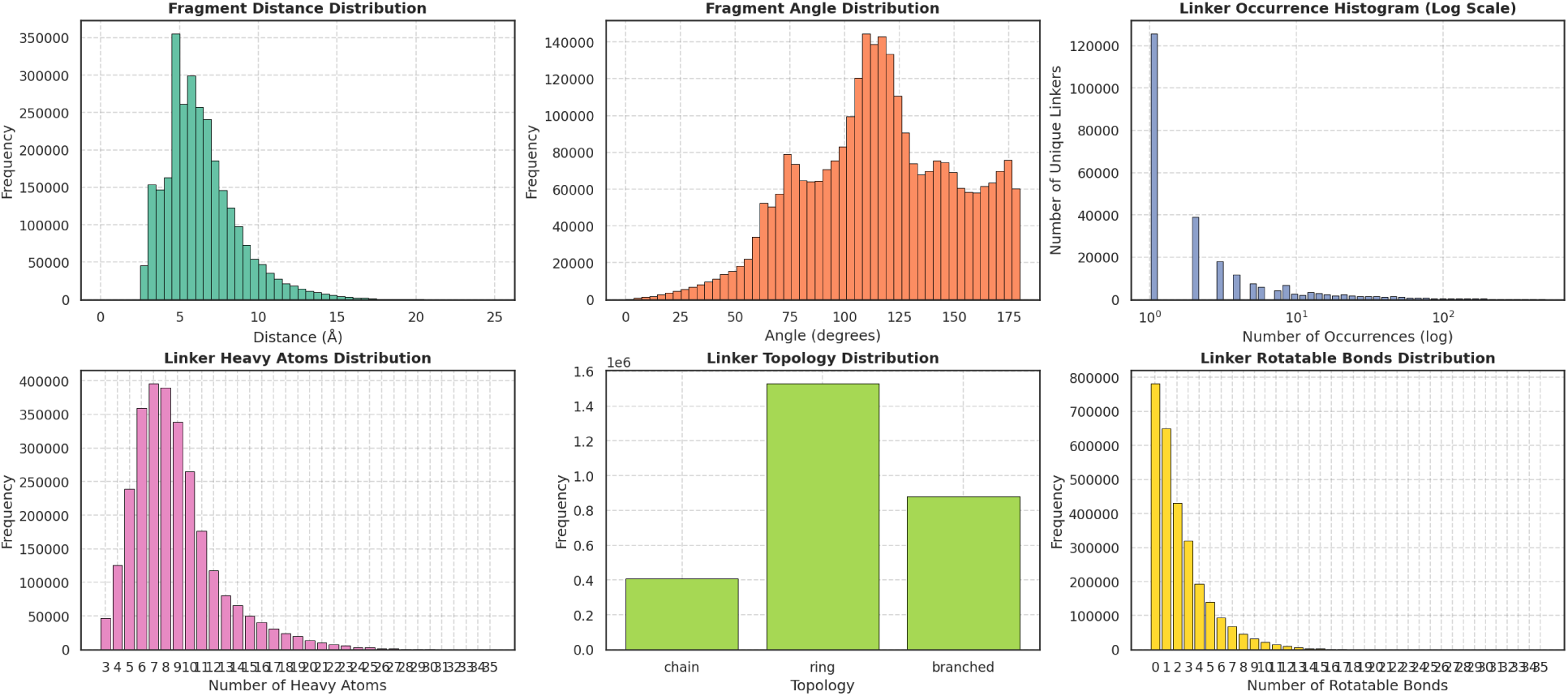
Fragment and linker property distributions for the hybrid balanced dataset (Hybrid), where the maximum occurrence of any unique linker is 500, with 10% reduction for linkers with occurences from 50 to 500. This intermediate strategy applies a softened capping mechanism, preserving the natural frequency hierarchy of common synthetic linkages while aggressively truncating the extreme occurrences seen in the full dataset. This balance ensures the model learns a diverse linker vocabulary without losing the inductive bias toward chemically intuitive, frequently synthesized motifs.

**Figure S3.5:**
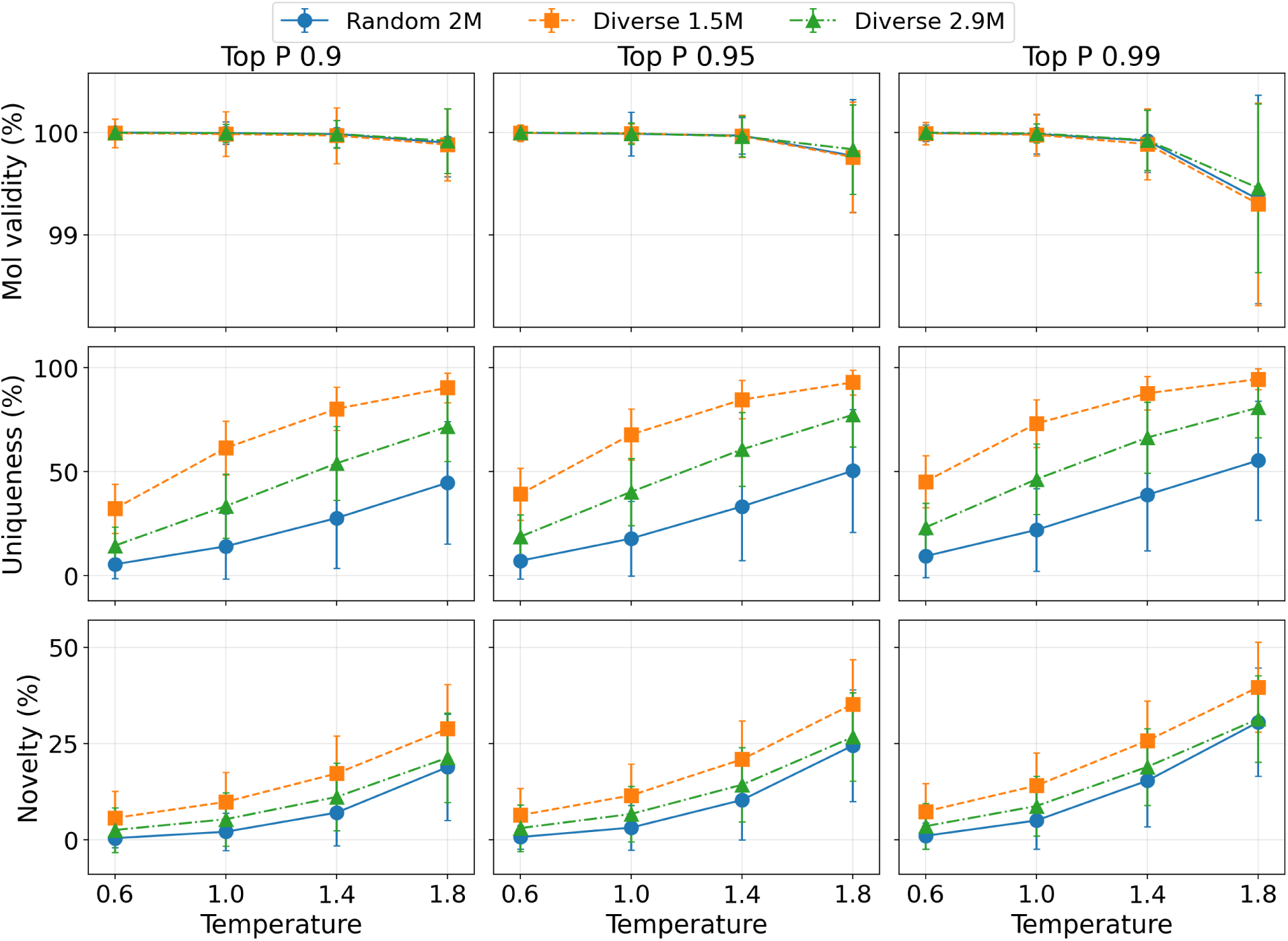
Model hyperparameter searching performances of unconditional generation on the ZINC-random dataset. The legend denotes the three evaluated models trained on different data splits: **Random 2M** (the long-tailed, unbalanced pool), **Diverse 1.5M** (the strict Cap50 rebalanced pool), and **Diverse 2.9M** (the hybrid-capped pool). Individual data points represent temperature settings (*T* ∈ {0.6, 1.0, 1.4, 1.8}), and the columns correspond to Top-P sampling values (top-*p* ∈ {0.9, 0.95, 0.99}). The rows display mean molecule validity (%), uniqueness among valid samples (%), and the novelty of valid linkers compared to the training linker set (%), with shared *y*-axes within each row. Error bars indicate the dispersion across instances and aggregated runs.

**Figure S3.6:**
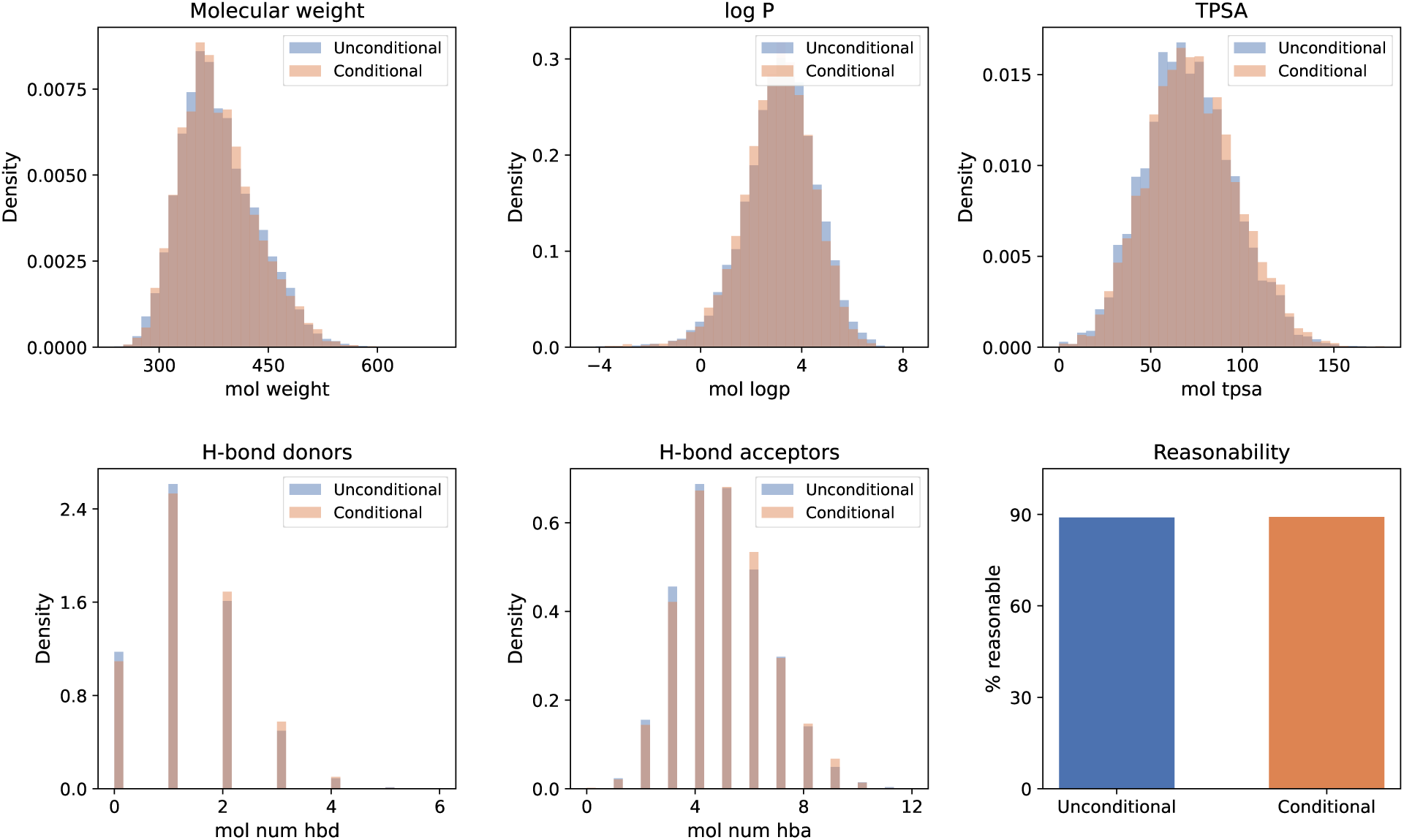
ZINC hard 1k, Ro5-focused prompt: empirical marginals for physicochemical properties compared to the LinkLlama unconditional baseline. The conditional generation (orange) has minimal changes compared to the unconditional baseline (blue) likely due to the fact that most training samples satify Ro5.

**Figure S3.7:**
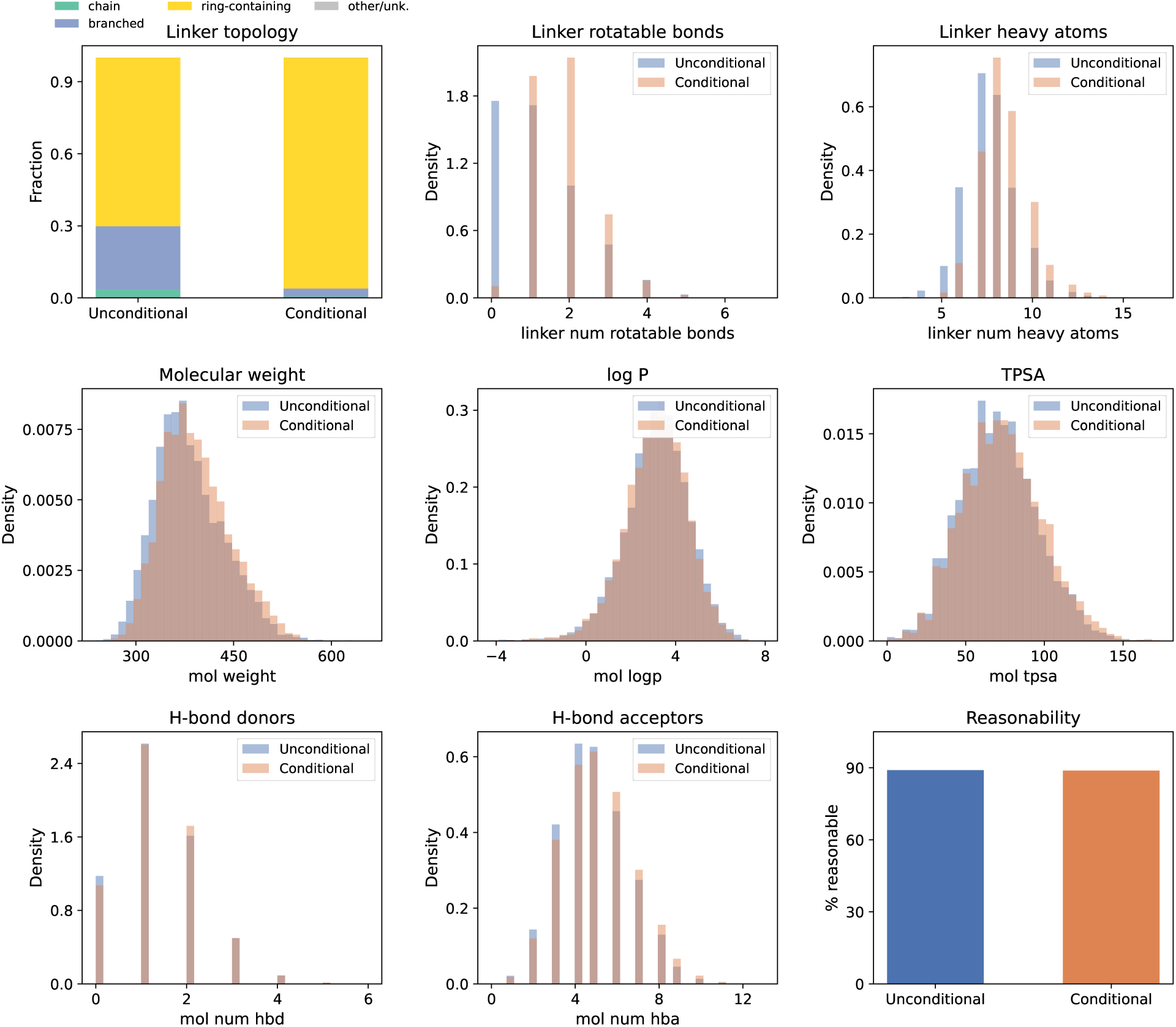
ZINC hard 1k, joint prompt (ring-containing linker, Ro5 bands, rotatable-bond and heavy-atom floors): empirical marginals for properties and linker topology compared to the unconditional baseline. The conditional generation (orange) demonstrates high prompt adherence across multiple simultaneous constraints, most notably forcing the linker topology to be nearly exclusively ring-containing, while shifting rotatable bonds (≥ 2) and heavy atoms (≥ 6) appropriately without violating Ro5 limits or degrading overall reasonability.

**Figure S3.8:**
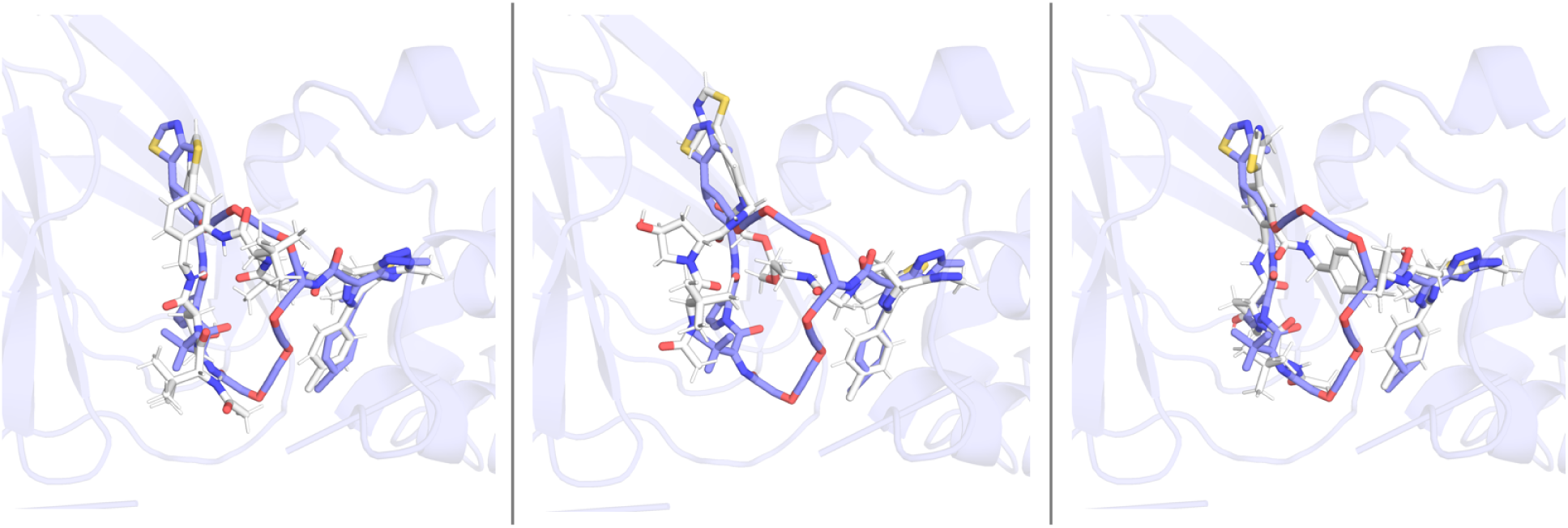
3D structural overlays of the docked poses of LinkLlama-generated linear PROTAC candidates on the crystal poses of the BRD4-VHL PROTAC ternary complex (PDB ID 6SIS). The reference macrocyclic PROTAC is shown in purple, while three top-performing generated linear candidates are depicted in white in the order of LinkLlama Resp 38, Resp 49, and Resp 82.

